# The Polar Night Shift: Annual Dynamics and Drivers of Microbial Community Structure in the Arctic Ocean

**DOI:** 10.1101/2021.04.08.436999

**Authors:** Matthias Wietz, Christina Bienhold, Katja Metfies, Sinhué Torres-Valdés, Wilken-Jon von Appen, Ian Salter, Antje Boetius

## Abstract

Change is a constant in the Arctic Ocean, with extreme seasonal differences in daylight, ice cover and temperature. The biodiversity and ecology of marine microbes across these extremes remain poorly understood. Here, using an array of autonomous samplers and sensors, we portray an annual cycle of microbial biodiversity, nutrient budgets and oceanography in the major biomes of the Fram Strait. In the ice-free West Spitsbergen Current, community turnover followed the solar cycle, with distinct separation of a productive summer state dominated by diatoms and carbohydrate-degrading bacteria, and a regenerative winter state dominated by heterotrophic Syndiniales, radiolarians, chemoautotrophic bacteria and archaea. Winter mixing of the water column replenishing nitrate, phosphate and silicate, and the onset of light were the major turning points. The summer succession of *Phaeocystis, Grammonema* and *Thalassiosira* coincided with ephemeral peaks of *Formosa, Polaribacter* and NS clades, indicating metabolic relationships between phytoplankton and bacteria. In the East Greenland Current, ice cover and greater sampling depth coincided with weaker seasonality, featuring weaker bloom/decay events and an ice-related winter microbiome. Low ice cover and advection of Atlantic Water coincided with diminished abundances of chemoautotrophic bacteria while *Phaeocystis* and *Flavobacteriaceae* increased, suggesting that Atlantification alters phytoplankton diversity and the biological carbon pump. Our findings promote the understanding of microbial seasonality in Arctic waters, illustrating the ecological importance of the polar night and providing an essential baseline of microbial dynamics in a region severely affected by climate change.

## INTRODUCTION

Microbes have fundamental roles in the marine biosphere and have been recognized as key components in global change biology^1,2^. Understanding the causes, complexity and consequences of microbial community dynamics requires continuous records in their environmental context. Time-series observations are beginning to discern the temporal variability and environmental drivers of marine microbiomes from diurnal to decadal scales^3,4^, but have mostly focused on temperate and tropical waters to date^5–13^. In contrast, temporal records from the polar oceans are rare. Pioneering studies have identified variable numbers, activities and community structures of polar microbes over time and space^14–21^, yet with limited temporal or spatial resolution.

Due to the extreme winter conditions, year-round sampling in polar waters requires autonomous observations, recently providing the first comprehensive annual records in the Arctic and Antarctic^22,23^. Such approaches can identify transition phases in the interplay between ocean physics and the ecosystem, for instance the onset of the spring bloom or the end of net-growth. In this regard, the polar night is of key interest, when physical mixing^24–26^ and microbial recycling of detrital and inorganic matter^27,28^ replenishes nutrients to fuel the subsequent phytoplankton bloom. Arctic phototrophic taxa are thought to persist in dormancy^29^, responding rapidly when light and stratification return^30^. Yet, microbial dynamics during the polar night within the oceanographic context remain virtually unknown.

Here, using an array of autonomous samplers and sensors, we portray microbial and oceanographic seasonality in the two major biomes of the Fram Strait. This main deep-water gateway to the central Arctic Ocean harbors the northward, permanently ice-free West Spitsbergen Current (WSC) and the southward, cold East Greenland Current (EGC) inside the marginal ice zone, with some recirculation in central Fram Strait (Fig. 1a). Our study is embedded in the long-term HAUSGARTEN and FRAM observatories studying primary productivity, benthopelagic coupling and deep-sea ecology since the 1990s^31,32^. The recent deployment of autonomous devices provides a key advance to characterize complete annual cycles, expanding on summertime snapshots of microbial diversity, biogeography and activity^33–39^. Annual records are also important to understand biological responses to the northward expansion of subarctic habitats, often coined Atlantification, which propagates through the entire food web^40,41^.

**Fig. 1.**
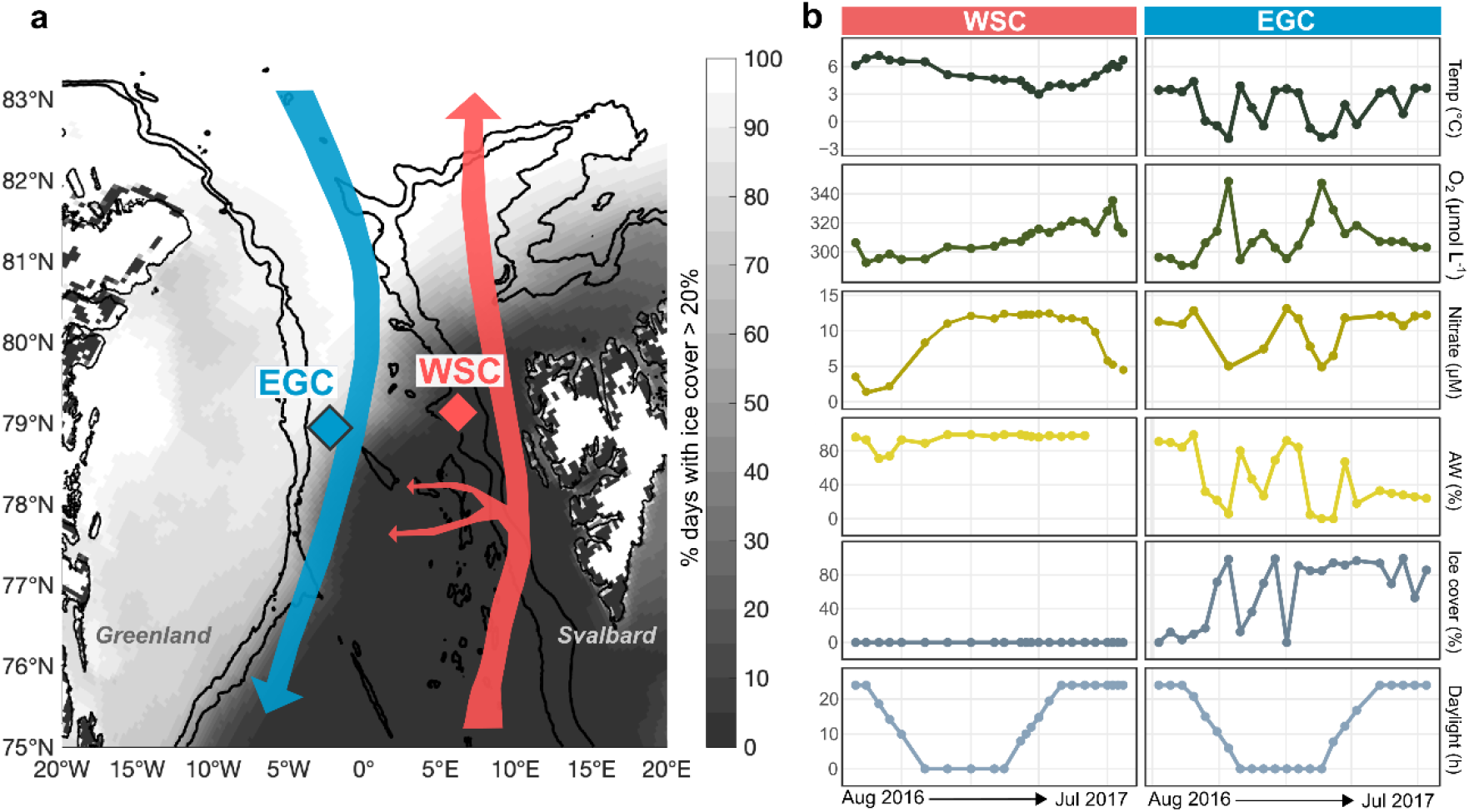
Study area and oceanographic conditions. **a:** Location of moored Remote Access Samplers in the East Greenland Current (EGC) and the West Spitsbergen Current (WSC) of Fram Strait, indicated in blue and red respectively. The small red arrows illustrate recirculation of Atlantic Water in central Fram Strait. The grayscale gradient illustrates the percentage of days with average ice cover of >20%. **b:** Annual water temperature (°C), concentrations of oxygen (µmol L^−1^) and nitrate (µM), the proportion of Atlantic Water (%), sea ice cover (%), and daylight hours.

Here we investigated how the influence of polar day and night shape seasonal shifts, expecting considerable differences between summer and winter microbiomes. We hypothesize that phototrophy- and heterotrophy-dominated periods in the ice-free WSC harbor markedly dissimilar microbial communities, while sea ice cover in the EGC favors winter-type communities year-round. This study improves the understanding of seasonal microbial biodiversity, the ecological importance of the polar night, and the effects of Atlantification in different areas of the Arctic Ocean. These insights are a key baseline to understand natural variability and human impact in a marine region under severe threat by climate change^42–45^, offering important perspectives on the present and future Arctic Ocean.

## RESULTS AND DISCUSSION

The present study elucidates microbial and oceanographic seasonality in the West Spitsbergen Current (WSC) and the East Greenland Current (EGC) of Fram Strait using automated, high-frequency sampling (Fig. 1a). For this purpose, seawater was autonomously collected and preserved *in situ* using moored Remote Access Samplers (RAS) in weekly to monthly intervals (Supplementary Table S1). In addition, oceanographic sensors continuously recorded temperature, salinity and oxygen concentration, informing about oceanographic conditions including the proportions of Atlantic Water (AW) and Polar Water (PW). After recovery, water samples were subjected to molecular fingerprinting of microbial communities and quantification of inorganic nutrients. Relative abundances of bacterial, archaeal and eukaryotic amplicon sequence variants (ASVs) were then evaluated in the oceanographic context, including satellite-derived sea ice and chlorophyll concentrations (Supplementary Table S1).

### Major annual dynamics and drivers

Environmental conditions and microbial community structure differed substantially between the two sampling sites (Fig. 1b, Extended Data Fig. 1−2). At the WSC mooring, ice-free AW prevailed throughout the year, with gradual physicochemical changes along the alternating stratification and mixing of the water column^46,47^. At the EGC mooring, deployed at the edge of the marginal ice zone, the intermittent advection of AW resulted in dynamic changes between polar (cold/ice-rich) and atlantified (warmer/low-ice) conditions (Fig. 1b). PW-dominated periods showed a typical physicochemical and microbial signature, whereas AW advection resulted in greater similarities to the WSC (Extended Data Fig. 1). This connection was strongest between AW proportions and bacterial composition (Extended Data Fig. 2; Spearman’s *rho* = 0.4, *p* = 0.00008). The EGC-RAS was deployed at 80 m depth to avoid collision with sea ice, i.e. located in the stratified halocline below the productive layer, compared to the WSC-RAS deployed at 25 m in the productive layer. Hence, differences between the WSC and EGC relate to hydrography and ice cover as well as sampling depth.

Nonetheless, the WSC and EGC shared a number of fundamental patterns in microbial diversity and community turnover. Microbial communities showed major compositional shifts over the annual cycle (Fig. 2a), with substantial month-to-month variability (PERMANOVA, *R*^*2*^ ≥ 0.6, *p* < 0.01) indicating dynamic microbiome structures year-round. With reference to the first sampling event, communities were most dissimilar around the March equinox before becoming more similar again towards peak polar day (Fig. 2b), illustrating light-driven temporal recurrence^48^. Notably, bacterial but not eukaryotic alpha-diversity correlated with daylight hours in both regions (Spearman’s *rho* = 0.6, *p* < 0.006).

**Fig. 2.**
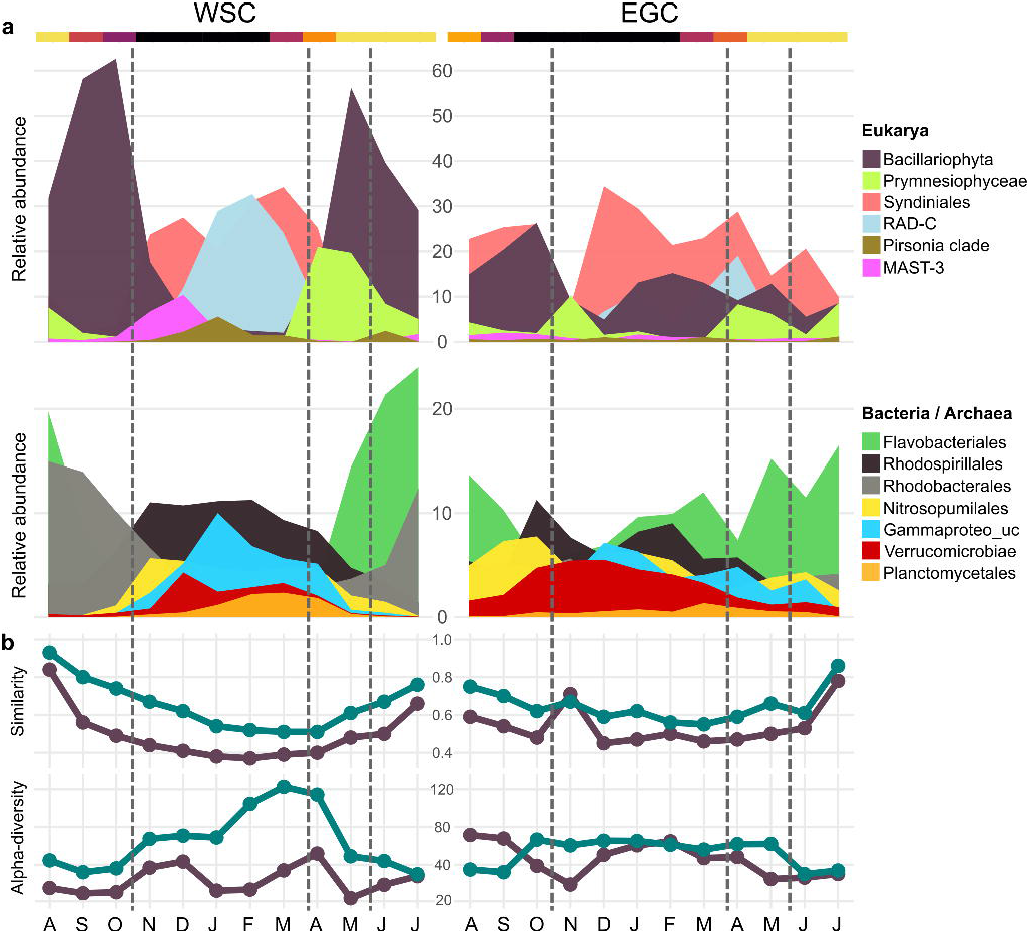
Annual cycle of microbial community structure and turnover. **a:** Relative sequence abundances (%) of eukaryotic, bacterial and archaeal taxa over the annual cycle. **b, upper panel:** Microbial community turnover (taxonomic similarities expressed as 1−Jenson-Shannon distance) compared to the first sampling event. **b, lower panel:** Microbial alpha-diversity (inverse Simpson index). Eukaryotes: purple; bacteria and archaea: green. Lines indicate the seasonal boundaries defined by multivariate evaluation of physicochemical and microbial dynamics (Fig. 3).

### Microbial and environmental seasonality

We contextualized major shifts and patterns in microbial and physicochemical variability (Figs. 2−4) to delineate the four seasons: spring (mid-April to mid-June), summer (mid-June to late-July), autumn (August to October) and winter (November to mid-April). Comparing all sampling events, community structures largely clustered by season and less by site (Extended Data Fig. 3). Hence, seasonality was overall a stronger driver than geographic distance, oceanographic differences and sampling depth. Each season showed a characteristic oceanographic and microbial signature (Figs. 3−4, Supplementary Table S2), supported by season-specific correlations between microbial taxa and environmental parameters (Extended Data Fig. 4). Winter microbiomes were most distinct, with up to ∼60% compositional dissimilarity to the other seasons. Greater seasonal contrasts in eukaryotic composition underlined the structuring role of light (Fig. 3b). In line with recent metagenomic evidence, these patterns indicate a considerable degree of temporal and functional specialization^49^.

**Fig. 3.**
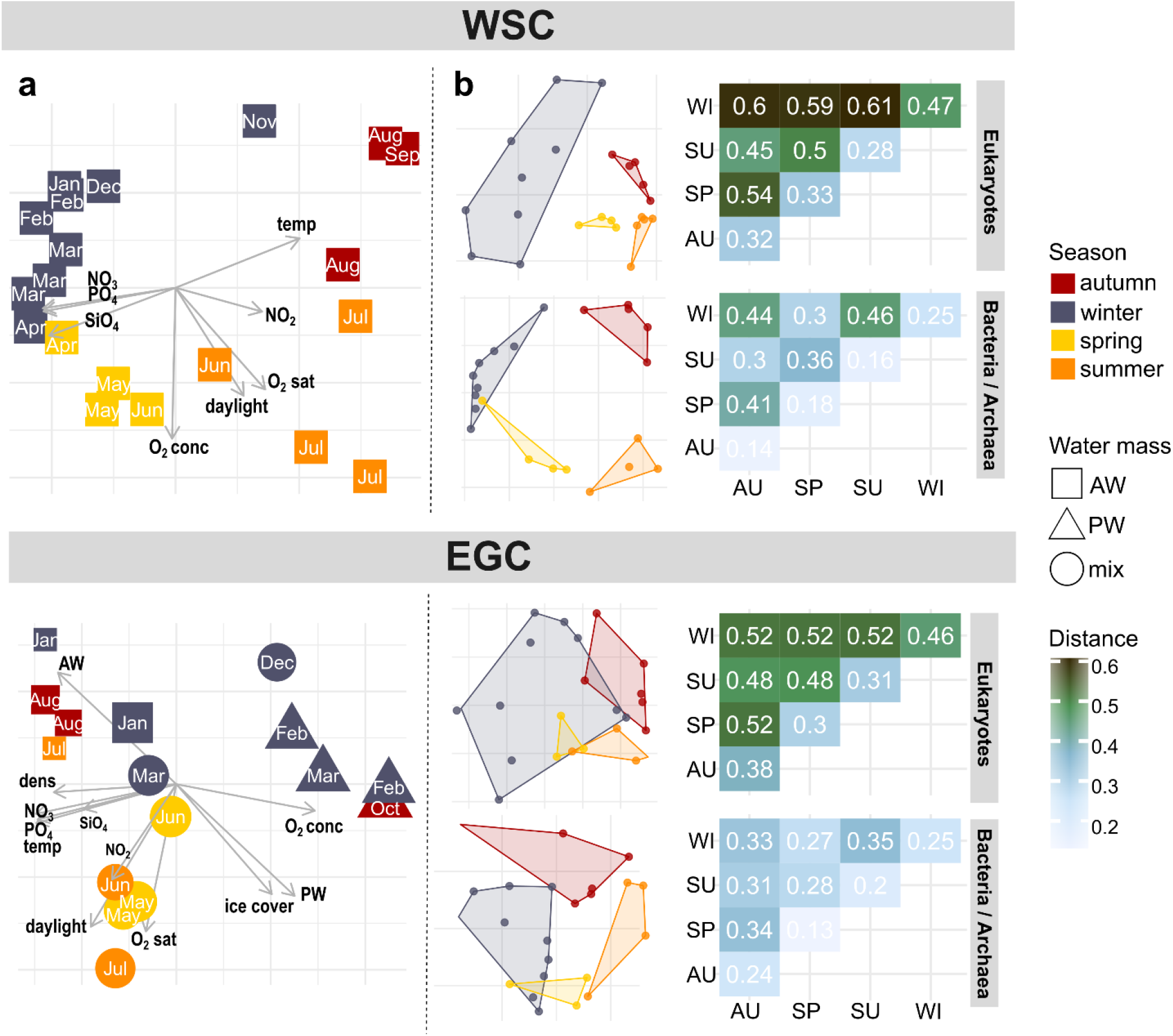
Microbial and environmental seasonality. **a:** Principal Component Analysis of environmental conditions. Components 1 and 2 explained 58/26% (WSC) and 60/14% (EGC), respectively, and hence the majority of physicochemical seasonality. For EGC, label size indicates percent ice cover. Only sampling events with complete environmental data were considered. **b:** Non-metric multidimensional scaling of Hellinger-transformed relative ASV abundances (stress values 0.06, 0.03, 0.12, 0.1 respectively) and corresponding Jensen-Shannon distances between and within seasons (larger numbers designating more dissimilar communities).

The WSC showed pronounced seasonality, both regarding environmental conditions (Fig. 3a) and microbial community composition (Fig. 3b). Daylight hours and temperature were the major drivers of eukaryotic variability (*R*^*2*^ *=* 0.2 respectively), whereas bacterial composition varied mostly with temperature (*R*^*2*^ = 0.4; PERMANOVA, *p* < 0.001) as previously observed in global TARA microbiomes^50^. Bacterial alpha-diversity peaked at the end of winter (Fig. 2b) when water temperatures were lowest (Fig. 1b), underlining the day-night shift as key transition event. ASVs associated with Bacillariophyta (i.e. diatoms) and Flavobacteriales predominated from spring to autumn (Fig. 2a), likely corresponding to metabolic interrelations through algal carbohydrates^51^. In contrast, heterotrophic eukaryotes (foremost Syndiniales and RAD-C radiolarians), archaea (Nitrosopumilales) and specific bacteria (e.g. Rhodospirillales) predominated in winter, with additional short-lived peaks of the diatom parasites *Pirsonia* and MAST (Fig. 2a). We consider these taxa as “microbial recyclers” persisting on detrital, inorganic or semi-refractory substrates^26,52^. The separation between photoautotrophy- and heterotrophy-driven periods of production and recycling were reflected in nutrient concentrations, with depletion in summer and replenishment during winter (Fig. 1b, Supplementary Table S1).

In the EGC, changes between polar and atlantified conditions happened on shorter time scales than seasons, causing a more variable community composition, turnover and diversity (Figs. 2–3). For instance, environmental conditions during AW advection in January resembled those in August (Figs. 1b, 3a). Daylight, temperature, hydrography and ice cover interrelated to comparable extents with microbial community structure (PERMANOVA, *R*^*2*^ = 0.1 for each factor, *p* < 0.05). Constant proportions of photoautotrophic and heterotrophic eukaryotes year-round, with ∼50% lower sequence abundances of diatoms than in the WSC (Fig. 2a, Extended Data Fig. 5), indicated a different food web structure. Low irradiance due to sea ice presumably repressed primary production at the surface, while simultaneously seeding diatoms and microbes into the underlying water (discussed in detail below). Furthermore, we observed a temporal lag in the occurrences of several taxa compared to the WSC. For instance, *Thioglobaceae* and SAR86 occurred in EGC-summer compared to WSC-spring; or *Mediophyceae* in EGC-autumn compared to WSC-summer (Fig. 4). Hence, microbial communities in the WSC and EGC can be variable across and within seasons, with temporal shifts resulting from later phytoplankton growth and the deeper RAS location. In the following, we present a detailed synopsis of seasonal patterns and specific events in chronological order from autumn 2016 to summer 2017.

**Fig. 4.**
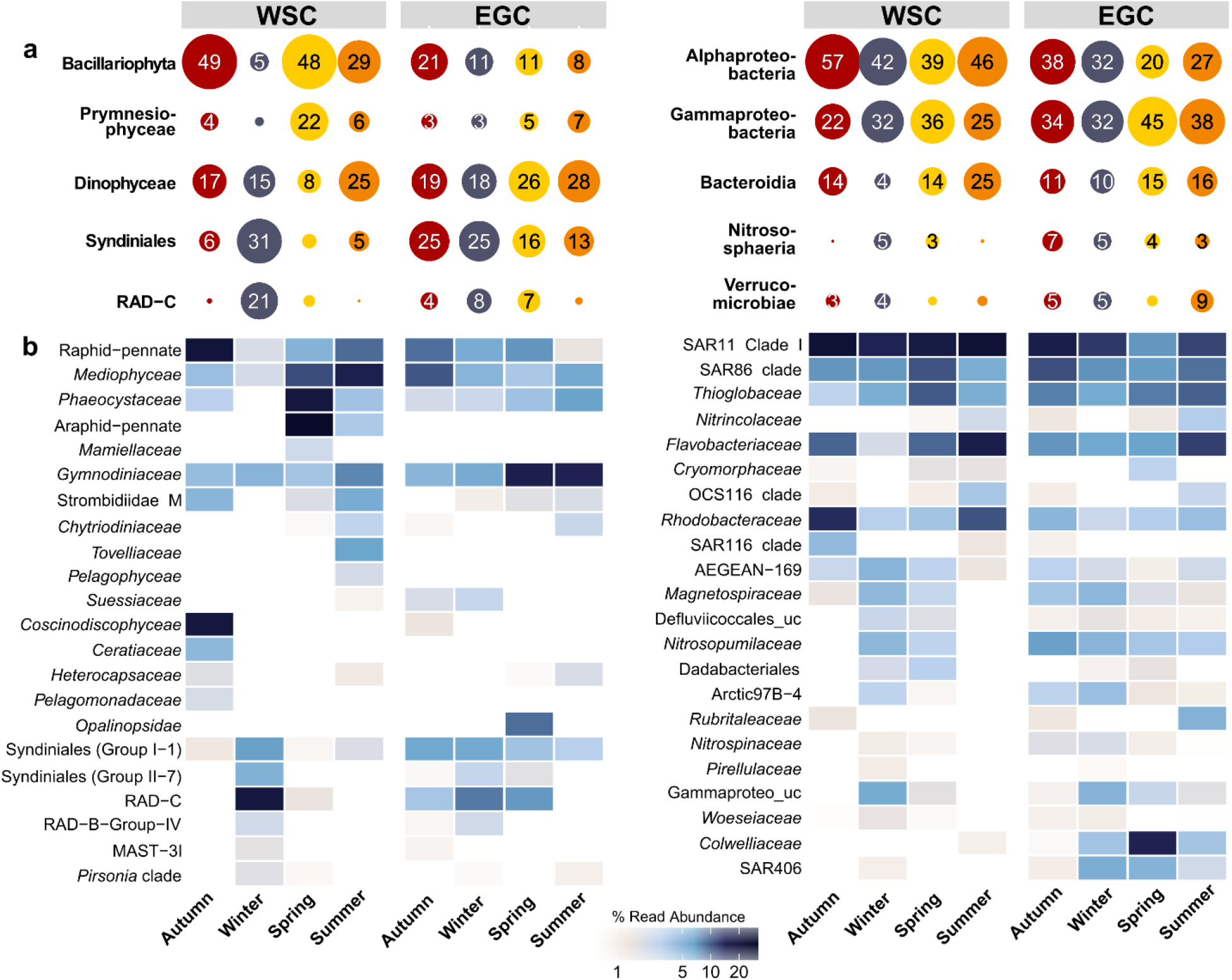
Microbes as indicators for seasons. Relative sequence abundances of major microbial classes (a) and families (b) by season. For classes only sequence abundances > 3% are shown.

### Autumn

Autumn in the WSC was characterized by nitrate, silicate and phosphate depletion and a specific community of *Coscinodiscophyceae, Ceratiaceae*, SAR116 and *Rhodobacteraceae* (Figs. 4−5, Supplementary Table S2). These patterns resemble a post-bloom state, with growing decay of summer phytoplankton^53^ and concurrent increase in mixotrophic dinoflagellates such as *Tripos*^54^. The prevalence of *Corethron, Rhizosolenia* and *Proboscia* sequences (Fig. 5b, Extended Data Fig. 5) matched microscopic cell counts in the Fram Strait^55^, corroborating our amplicon-based results. Similar autumn patterns in the Southern Ocean indicate bi-polar seasonal preferences of *Coscinodiscophyceae* diatoms, likely facilitated by their ability to overcome silicate limitation^56^, use ammonium instead of nitrate^22^, and resist grazing^57^. Appearance of chytrid fungi and fungi-like *Labyrinthulaceae* at maximal nutrient depletion in October (Extended Data Fig. 6) indicates saprophytic activity on decaying algae^58–60^. Among bacteria, up to 13-fold higher abundances of *Cand*. Puniceispirillum and other SAR116 members as well as the *Rhodobacteraceae* taxa *Ascidiaceihabitans, Amylibacter and Planktomarina* (Fig. 5b) were probably fueled by DMSP and senescence compounds from decaying phytoplankton^61–63^. Detection of *Luteolibacter* from the *Rubritaleaceae* family (Fig. 5b) suggested concurrent particle formation, typical processes in ageing phytoplankton^64^ as observed in coastal Svalbard^65^.

**Fig. 5.**
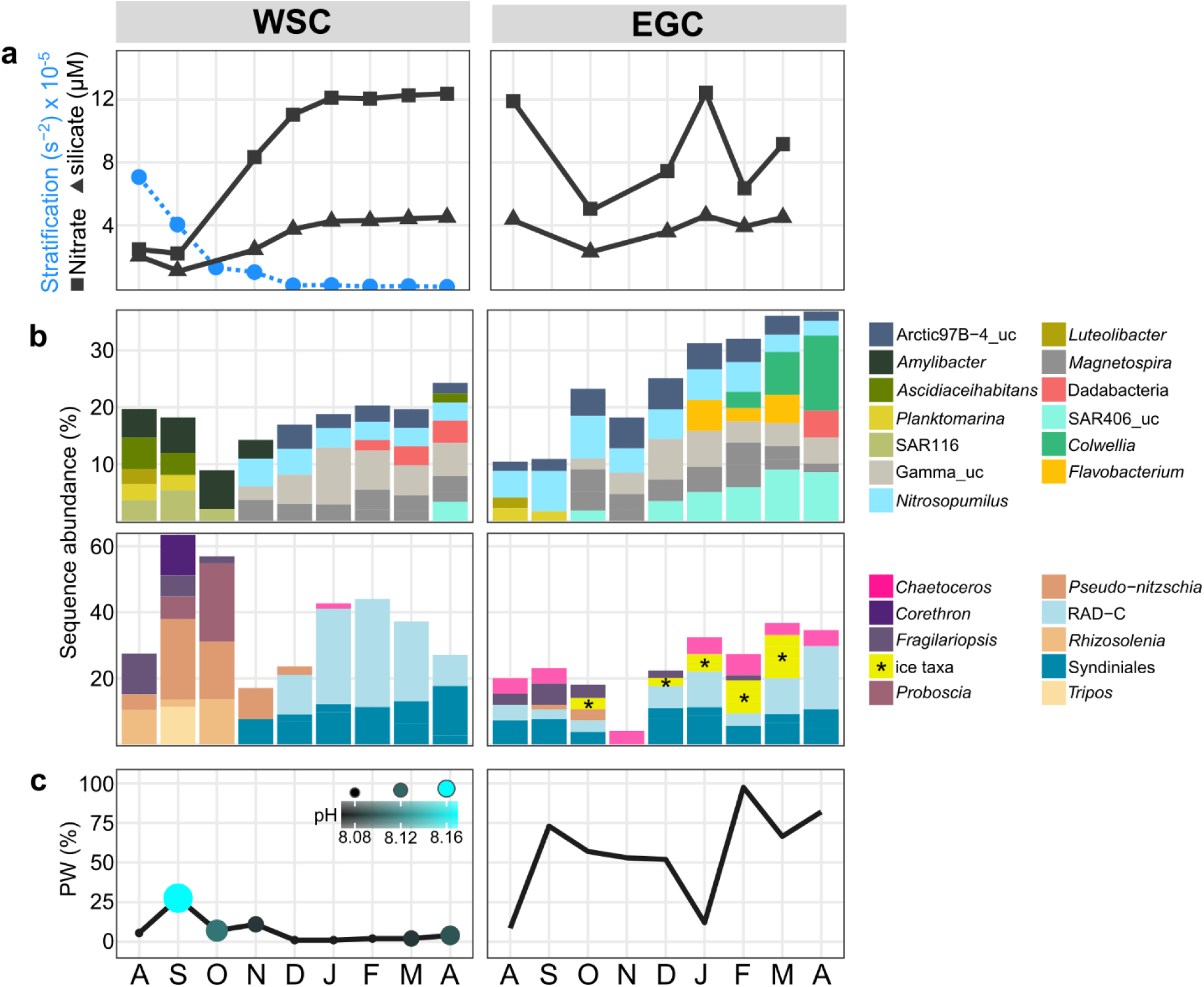
Autumn and winter. **a:** Concentrations of nitrate (squares) and silicate (triangles) in relation to stratification (blue; only available for the WSC). **b:** Microbial genera with increased proportions in autumn or winter. “Winter-ice” eukaryotes are combined (marked with asterisks; see Extended Data Fig. 7a for abundances of each genus). **c:** pH values (only available for the WSC) and proportions of PW.

*Fragilariopsis* co-occurred in the WSC and the EGC during early autumn (Fig. 5b). We hypothesize that this typically ice-associated taxon was transported to the WSC by advection, considering the higher proportion of PW during this time (Fig. 5c). This event also covaried with higher pH, with potential metabolic effects on prevalent taxa such as *Pseudo-nitzschia*^66,67^. Otherwise, the EGC displayed quite different dynamics. Peaking diatom abundances characterized autumn as major photosynthetic period (Fig. 4, Extended Data Fig. 5), featuring SAR86 characteristic of WSC-spring (Fig. 4). We attribute this late bloom to the minimum ice cover in autumn (Fig. 1b, Supplementary Table S2), enhancing light penetration, melt-induced stratification and ice seeding^68,69^.

### Winter

The WSC and EGC shared elevated abundances of *Magnetospiraceae, Nitrospinaceae*, the Arctic97B-4 clade and unclassified Gammaproteobacteria (Figs. 4, 5b), although winter-summer contrasts of these taxa were stronger in the WSC (average Kruskal-Wallis significance *p* ≤ 0.003 vs. 0.02 in the EGC). Furthermore, Dadabacteriales co-occurred from February (WSC) or late March (EGC) and might contribute to the recycling of microbial and other organic matter^70^. Fundamental regional differences were the heterotrophic “recycling state” in the WSC compared to ice-related microbial signatures in the EGC.

#### Heterotrophic winter communities of the WSC

The increase of Syndiniales, parasitic recyclers of phytoplankton biomass^71–73^, in November marked the onset of winter. Bacterial diversification and nutrient replenishment (Figs. 2, 5, Extended Data Fig. 6) followed the breakdown of summer stratification^74,75^, with highest mixing of the water column in January (Fig. 5a). At this time, heterotrophic eukaryotes constituted up to ∼70% of sequences and nutrient standing stocks were restored (Fig. 2a, 5a). The parallel decline of phototrophs to a combined relative abundance of <5% (Extended Data Fig. 5) indicated complete mixing as one central turning point of the annual cycle, emphasizing the ecological importance of Arctic winter^24^. Notably, this also illustrates that only a small “seed bank” overwintered to initiate the following spring bloom. The upward transport of microbes during mixing likely enriched the metabolic potential^76,77^. For instance, appearance of deep-water RAD radiolarians^78^ possibly contributed to the recycling of phytoplankton biomass. Stratification potentially also influenced the temporal succession of different Syndiniales clades over winter (Extended Data Fig. 6).

The co-occurrence of *Nitrosopumilaceae* and *Nitrospinaceae* (Figs. 4, 5b), the major drivers of marine nitrification, suggests an interactive niche with initial oxidation of ammonia or urea by *Nitrosopumilaceae* and subsequent nitrite oxidation by *Nitrospinaceae*^79–81^. In addition, the *Magnetospiraceae* family (Rhodospirillales) might recycle nitrogen by fixation and contribute to a yet underestimated nitrogen source^82,83^, although no related nitrogenase sequences were yet detected in the Arctic^84^. Furthermore, metaproteomic data indicate that *Magnetospiraceae* also perform CO_2_ fixation and thiosulfate oxidation^82,85,86^. Genomic and metabolic evidence from other Arctic and Antarctic winter microbiomes suggests consistent roles of *Nitrosopumilaceae, Nitrospinaceae* and *Magnetospiraceae* during winter at both poles^19,27,87–93^. Further potential recyclers are the *Pirellulaceae* and *Woeseiaceae* through ammonia oxidation and denitrification respectively^94,95^. The winter niche of the poorly described *Defluviicoccales* clade is potentially fueled by stored glycogen^96,97^ or unsaturated aliphatics^98^. Overall, the observed elevated abundances of diverse heterotrophic and chemoautotrophic taxa highlight the polar night as an important recycling phase before the spring bloom. Furthermore, the polar night microbiome is not static, but responsive to certain stimuli such as mixing.

#### The microbial winter loop in the EGC

Unique to the EGC was the persistence of raphid-pennate diatoms and flavobacteria throughout winter (Fig. 4), contrasting their light-controlled seasonality in the WSC. We attribute these signals to ice melt and repeated seeding events, following intermittent water temperatures of >2°C during AW advection in January (Fig. 1b). The diatoms *Bacillaria* and *Naviculales*, together with *Polarella* flagellates and *Chrysophyceae*, constituted up to 15% of sequences after the ice melt between February and March (Fig. 5b, Extended Data Fig. 7a). All of these taxa occur in sea ice and the underlying water^60,99–102^, suggesting an ice-dependent heterotrophic food web. Ice algae produce copious amounts of storage polysaccharides and extracellular polymeric substances, fueling bacterial growth in the underlying water^69,103^. *Bacillaria* exudates are a valuable nutrient source for bacteria^104^, as is chrysolaminarin from diatoms and *Chrysophyceae*^105,106^. The flavobacterial winter assemblage largely corresponded to a *Flavobacterium* ASV constituting ∼10% between January and March (Fig. 5b, Extended Data Fig. 7a). This ASV shared >99% 16S rRNA similarity with *Flavobacterium frigidarium*, a psychrophilic genus with laminarinolytic abilities^107^. Detection of related sequences on ice-algal aggregates^60^ supports a presumed niche through utilization of ice-algal carbohydrates. Overall, these light-independent, ice-fueled processes might explain signatures and activities of specific microbial taxa in other Arctic regions under warming^108–110^.

An EGC-specific winter taxon was the SAR406 clade, peaking at 9% relative sequence abundance in March and remaining detectable until summer. In addition, the frequently ice-associated genus *Colwellia* increased from February until a major bloom of >20% in mid-June (Figs. 5b, 6a). We propose that the average ice cover of ∼60% between February and July (Supplementary Table S2) sustained these winter taxa. Both SAR406 and *Colwellia* markedly correlated with ice cover (Spearman’s *rho* = 0.7, *p* < 0.0004), highlighting the role of sea ice for microbial biodiversity in polar regions. SAR406 is also abundant and active below the Antarctic shelf ice^93^ and might participate in sulfur cycling^111^, including DMSP from ice algae^112^. Increasing loss of sea ice might therefore diminish the recycling of inorganic substrates.

### Spring and summer

#### Microbial succession in the WSC

Once daylight reached ∼20h in mid-April, the microbial system quickly returned to a phototrophic state. The winter-spring transition occurred within two weeks and hence as rapid as in warmer Pacific waters^113^, emphasizing light as the fundamental trigger. Eukaryotic composition changed ahead of bacterial communitieswhereas heterotrophs responded with delay of approximately 14 days to the prime photosynthetic peak (Extended Data Fig. 6). We observed three distinct bloom stages, featuring phototrophic pioneers *(Phaeocystis* and *Chaetoceros*) followed by araphid-pennate diatoms (*Grammonema*) and centric diatoms (*Thalassiosira*). A comparable three-stage summer bloom in the preceding year in nearby Kongsfjorden^21^ suggests recurrent principles in the ice-free Fram Strait. The swift replacement of eukaryotic heterotrophs by photoautotrophs (Fig. 3b, Extended Data Fig. 6) suggests considerable energy fluxes around the winter-spring transition, corresponding to transparent exopolymer particles^36^ and sinking radiolarians^114^, with effects on zooplankton grazing and deep-water biology^115,116^. The two early bacterial responders *Aurantivirga* and SAR92 (Extended Data Fig. 6) also appeared first in the Antarctic spring bloom^22^, indicating comparable ecological roles. The *Grammonema* abundance of >50% in May coincided with peaking chlorophyll (Fig. 6a), indicating substantial biomass and release of carbohydrates^117^. In this context, intermittent peaks of *Formosa, Polaribacter* and NS clades (Fig. 6a) illustrate carbohydrate-driven niches, resembling diatom-flavobacteria relationships in temperate and Antarctic waters^51,118–120^.

**Fig. 6.**
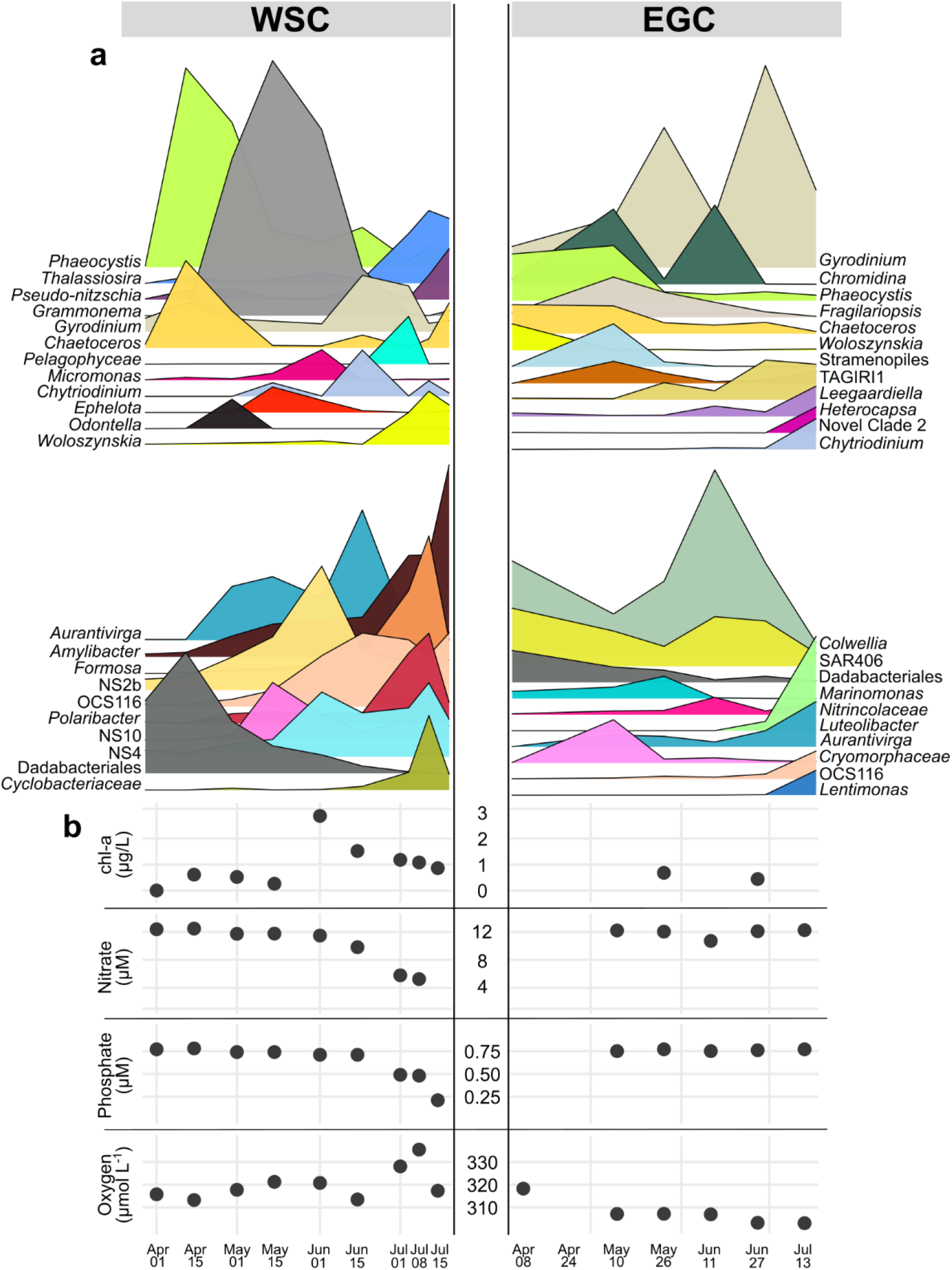
Spring and summer. **a:** Relative abundances of dominant eukaryotic and bacterial genera. **b:** Concentrations of chlorophyll, nitrate, phosphate and oxygen.

*Thalassiosira* was specific for summer and the final bloom stage, when nitrate and phosphate started declining and oxygen concentrations were highest (Fig. 6). The relative increase of mixotrophic flagellates (e.g. *Gyrodinium* and *Woloszynskia*) and concurrently decreasing chlorophyll indicates a shifting trophic structure, with probable effects on zooplankton activities^121–123^. Increase of the roseobacter *Amylibacter* (formerly NAC11-7) to 15% relative abundance emphasized the beginning transition to the autumn post-bloom where *Rhodobacteraceae* dominated (Fig. 4). Accordingly, the final sampling event in July 2017 clustered with August 2016 but not with the other summer samples (Fig. 3a). We hypothesize concurrent generation of detritus particles, given the typical termination of diatom blooms by aggregation^64,124^ and the association of *Amylibacter* with related particles^125^. Furthermore, the appearance of suctorian ciliates (*Ephelota*) and ectoparasitoid dinoflagellates (*Chytriodinium*) indicates the start of parasitism on diatoms and larger metazoans^126,127^.

#### Absence of major phototrophic peaks in the EGC

Sequence abundances of *Phaeocystis* only reached 9% compared to 40% in the WSC, presumably forming an under-ice bloom^30^ as ice cover was ∼90% at this time (Fig. 1b). Diatom abundances resembled those during winter (Extended Data Fig. 5), and chlorophyll concentrations were threefold lower than the WSC peak (Fig. 6b). *Fragilariopsis* and *Chaetoceros* together only constituted <10% of eukaryotic sequences, although nutrients were not limiting (Fig. 6b, Supplementary Table S2). Instead, Syndiniales prevailed over summer, with additional peaks of heterotrophic stramenopile and TAGIRI lineages (Figs. 4, 6a). The flagellates *Chromidina* (Ciliophora: Opalinopsidae) and *Gyrodinium* (Dinoflagellata: Gymnodiniaceae) constituted between 13 and 35% of eukaryote sequences depending on ice cover. *Chromidina* was specific to spring and is normally considered as animal parasite, suggesting yet undescribed niches in the marginal ice zone potentially related to the concurrent occurrence of amphipods^128^. The lower extent of primary production coincided with lower flavobacterial abundances compared to the WSC (Figs. 2a, 4b), including delayed appearance of the typical phytoplankton associates OCS116, *Lentimonas* and *Luteolibacter*^129,130^ (Fig. 6a). Peaks of gammaproteobacterial Oceanospirillales (*Nitrincolaceae, Marinomonas*) and an EGC-specific *Cryomorphaceae* lineage (Fig. 6a) supported the notion of a different food web based on other substrates. The presence of ice cover over summer suggests a continuous input of ice-derived substrates. These likely fueled the major peak of *Colwellia*, which can rapidly and efficiently grow on both high- and low-molecular weight organic matter from sea ice^69^.

## ECOLOGICAL CONCLUSIONS

Our comprehensive analysis of microbial seasonality in the Fram Strait by autonomous year-round sampling identified marked seasonal contrasts, distinct transition events, and short-term dynamics during mixing events and other environmental fluctuations^11^. The characterization of bloom stages, ephemeral abundance peaks and polar night characteristics promotes the understanding of the drivers and timescales of microbial seasonality in ice-covered and ice-free waters. These insights yield a number of fundamental ecological conclusions, with implications for the current and future Arctic Ocean.

1. We identified major principles of seasonality in the Arctic Ocean, with different levels of variability: (i) marked seasonal contrasts especially in the ice-free WSC; (ii) month-to-month dynamics following temperature and stratification gradients in the WSC; and (iii) ephemeral peaks related to diatom-flavobacterial relationships (WSC) or to intermittent changes in ice cover and hydrography (EGC).
2. Dynamics in the WSC represent key principles of seasonality in the ice-free, pelagic Arctic Ocean: *Phaeocystis* as daylight pioneer followed by araphid-pennate diatoms and maximum chlorophyll (spring); declining nitrate and concurrent shift towards mixotrophic flagellates and centric diatoms (summer); minimum nutrients and highest temperatures when *Coscinodiscophyceae* diatoms and oligotrophic bacteria prevailed (autumn); and heterotrophic microbial recyclers and nutrient replenishment during vertical mixing (winter). Comparable observations have been made in a year-round study using Niskin-based sampling^19^, illustrating that autonomous techniques provide results consistent with traditional approaches while considerably increasing temporal resolution. The present study has remarkable similarities to a RAS-based study in the open Southern Ocean, which reports *Coscinodiscophyceae* in autumn, *Aurantivirga* and SAR92 as first bacterial responders, and *Amylibacter* at the summer-autumn transition^22^. This suggests fundamental “bi-polar” patterns of microbial seasonality, only discernable by autonomous sampling.
3. The EGC mooring served to analyze dynamic effects of the extent of ice cover: it (i) extended the duration and abundance of winter taxa such as SAR406 and *Colwellia* when ice remained high in summer, (ii) light-limited phytoplankton growth, which only increased at low ice in autumn, and (iii) sustained a distinct heterotrophic signature throughout the year. Increasing light availability at low ice cover favored *Phaeocystis, Thalassiosira*, OCS116 and *Aurantivirga* (Extended Data Fig. 7b). These dynamics are sentinels of how the future EGC might shift from an ice-controlled to a light-controlled habitat^34,131^; affecting the fate of blooms^132–134^, the quality and quantity of algae-derived organic matter^135^, and the biological carbon pump^136^. Elevated photosynthesis and higher organic load might stimulate the microbial loop^137,138^, with rapid remineralization of ice-derived organic matter^25,139,140^ at the expense of autotrophic sulfur and nitrogen metabolism^111,141^.
4. Atlantification of the Arctic may enhance early blooms of *Phaeocystis*^30,33,142^ and substantially alter biogeochemical fluxes, considering the associated production of TEP that serves as microbial substrate, microhabitat and downward vehicle of organic matter. If stratification becomes stronger and more permanent with increasing temperatures, wintertime convection could diminish and deep-water “recycling taxa” disappear from the winter assemblage, with yet unknown ecological consequences.

In conclusion, the demonstrated seasonal dynamics and drivers of Arctic Ocean microbiomes are essential to understanding ecosystem functioning over polar day and night. Considering the anticipated impact of climate change on polar regions and the relevance of microbes in future ocean scenarios^1,2^, our evidence contributes to assessing natural variability, anthropogenic impact and microbial dynamics of the warming Arctic Ocean.

## MATERIALS AND METHODS

### Sampling approach

Within the framework of the FRAM long-term marine observatory (https://www.awi.de/en/expedition/observatories/ocean-fram.html), Remote Access Samplers (McLane, East Falmouth, MA) were deployed in July 2016 on seafloor moorings F4-S-1 in the core WSC (79.0118N 6.9648E) and EGC-3 in the marginal ice zone (78.831 N −2.7938E) at 20 and 80 m depth respectively (Fig. 1a). The fixed-point (Eulerian) approach, established standard in biological time-series observations, includes variation relating to the continuous movement of water masses. These also moved the RAS up and down in the water column, resulting in variable actual sampling depths (on average 40 m for F4-S-1 and 90 m for EGC-3; Supplementary Table S1). RAS frames were equipped with 48 sterile sampling bags, each containing 700 µL saturated mercuric chloride solution. At each programmed sampling event, two water samples of 500 mL were autonomously pumped an hour apart into individual sampling bags and fixed by the mixing with mercuric chloride. RAS were recovered in August 2017 and water samples immediately filtered through 0.22µm Sterivex filter cartridges (Millipore, Burlington, MA). Filters were frozen at –20°C until DNA extraction in the home lab.

### DNA extraction and amplicon sequencing

DNA was extracted using the PowerWater kit (QIAGEN, Germany) according to the manufacturer’s instructions, and quantified using Quantus (Promega, Madison, WI). Microbial 16S and 18S rRNA gene fragments were amplified using primers 515F–926R^143^ and 528iF–964iR^38^ respectively. Libraries were prepared according to the 16S Metagenomic Sequencing Library Preparation protocol (Illumina, San Diego, CA). 16S and 18S rRNA gene fragments were sequenced using MiSeq technology at CeBiTec (Bielefeld, Germany) or Alfred Wegener Institute respectively (Supplementary Methods).

### Sequence analysis

After primer removal using cutadapt^144^ reads were processed into amplicon sequence variants (ASVs) using DADA2 v1.14.1^145^. For 16S amplicons, reads from two independent IIlumina runs were merged after error learning as per the author’s recommendation. Reads were quality-trimmed based on DADA quality profiles. After singleton removal, we obtained on average 62,000 16S rRNA and 99,000 18S rRNA reads per sample (Supplementary Table S3) that sufficiently covered community composition (Extended Data Fig. 8). Prokaryotic and eukaryotic reads were taxonomically classified using Silva v138 and PR^2^ v4.12 databases respectively (Supplementary Methods). Subsequently, two samples collected at >200m depth were discarded to omit potential deep-water signatures.

### Mooring and satellite data

Temperature, salinity, oxygen concentration and oxygen saturation were derived from a CTD sensor attached next to the RAS, confirming consistent properties of the two water samples from each date. The physical sensors were manufacturer-calibrated and processed in accordance with https://epic.awi.de/id/eprint/43137. Raw and processed mooring data are available at^146^. For chemical sensors, the raw sensor readouts are reported. Partial CO_2_ pressure and pH were measured in the WSC only. Water masses and the fraction of Atlantic and Polar Water were characterized following Richter and colleagues^147^ at each sampling event (Supplementary Methods). Sea ice concentrations derived from the Advanced Microwave Scanning Radiometer sensor AMSR-2^148^ were downloaded from Institute of Environmental Physics, University of Bremen (https://seaice.uni-bremen.de/sea-ice-concentration-amsr-eamsr2). Surface chlorophyll concentrations measured with the Sentinel 3A OLCI were downloaded from https://earth.esa.int/web/sentinel/sentinel-data-access. For all satellite-derived data, we considered grid points within a radius of 15 km around the moorings.

### Nutrient quantification

Analyses were done using a QuAAtro Seal Analytical segmented continuous flow autoanalyser following standard colorimetric techniques. The accuracy of the analysis was evaluated using KANSO LTD Japan Certified Reference Materials, with corrections applied as required. After quality control in comparison with reference data, samples with QC score ≥4 were excluded from further analyses (labelled NA in Supplementary Table S1).

### Statistical evaluation

Data analysis was done in R v3.6.1^149^ implemented in RStudio (https://rstudio.com). In short, alpha-diversity (inverse Simpson index) and rarefaction curves were computed on raw ASV counts, excluding metazoan, chloroplast and mitochondrial sequences. Subsequently, we only considered reads with ≥3 counts in ≥2 samples. NMDS was performed using Jenson-Shannon distances on Hellinger-transformed ASV counts. Seasons were defined based on multivariate patterning of oceanographic parameters and microbial community composition (Figs. 2−3). Water masses were categorized as AW or PW at proportions of >80% respectively; and 20–80% as mixture of both. Statistical differences were computed by PERMANOVA, Wilcoxon rank-sum test, or Kruskal-Wallis plus Bonferroni-corrected Dunn’s post-hoc test whenever appropriate. Pairwise associations were assessed by Spearman’s correlation coefficients.

### Data availability

Amplicon sequences have been deposited in the European Nucleotide Archive under accession numbers PRJEB43890 (16S) and PRJEB43504 (18S) using the data brokerage service of the German Federation for Biological Data (GFBio) in compliance with MIxS standards. Environmental data has been deposited at PANGAEA (see Supplementary Table S1). Code and input files for reproducing workflow and figures are available at https://github.com/matthiaswietz/RAS-1617.

## Acknowledgements

We thank Jana Bäger, Theresa Hargesheimer, Daniel Scholz, Rafael Stiens and Lili Hufnagel for RAS operation; Normen Lochthofen, Janine Ludszuweit, Lennard Frommhold and Jonas Hagemann for mooring operation; Jakob Barz and Swantje Rogge for DNA extraction and library preparation; Halina Tegetmeyer for quality-control and sequencing of 16S rRNA amplicons; and Laura Wischnewski for nutrient analysis. Christiane Hassenrück, Stefan Neuhaus, Pier L. Buttigieg, Magda Cardozo and Andrew B. Collier contributed bioinformatic assistance. We thank Eva-Maria Nöthig for constructive discussions and the entire FRAM team for excellent collaboration at home and sea. The captain, crew and scientists of RV Polarstern cruises PS99 and PS107 are gratefully acknowledged. Ship time was provided under grants AWI_PS99_00 and AWI_PS107_05. This project has received funding from the European Research Council (ERC) under the European Union’s Seventh Framework Program (FP7/2007-2013) research project ABYSS (Grant Agreement no. 294757) to AB. Additional funding came from the Helmholtz Association, specifically for the FRAM infrastructure and from the Max Planck Society. This publication is Eprint ID XXXXX of the Alfred Wegener Institute Helmholtz Center for Polar and Marine Research, Bremerhaven, Germany.

