## Supplementary figures and images for "The Polar Night Shift: Annual Dynamics and Drivers of Microbial Community Structure in the Arctic Ocean"

### Extended Data Fig. 1

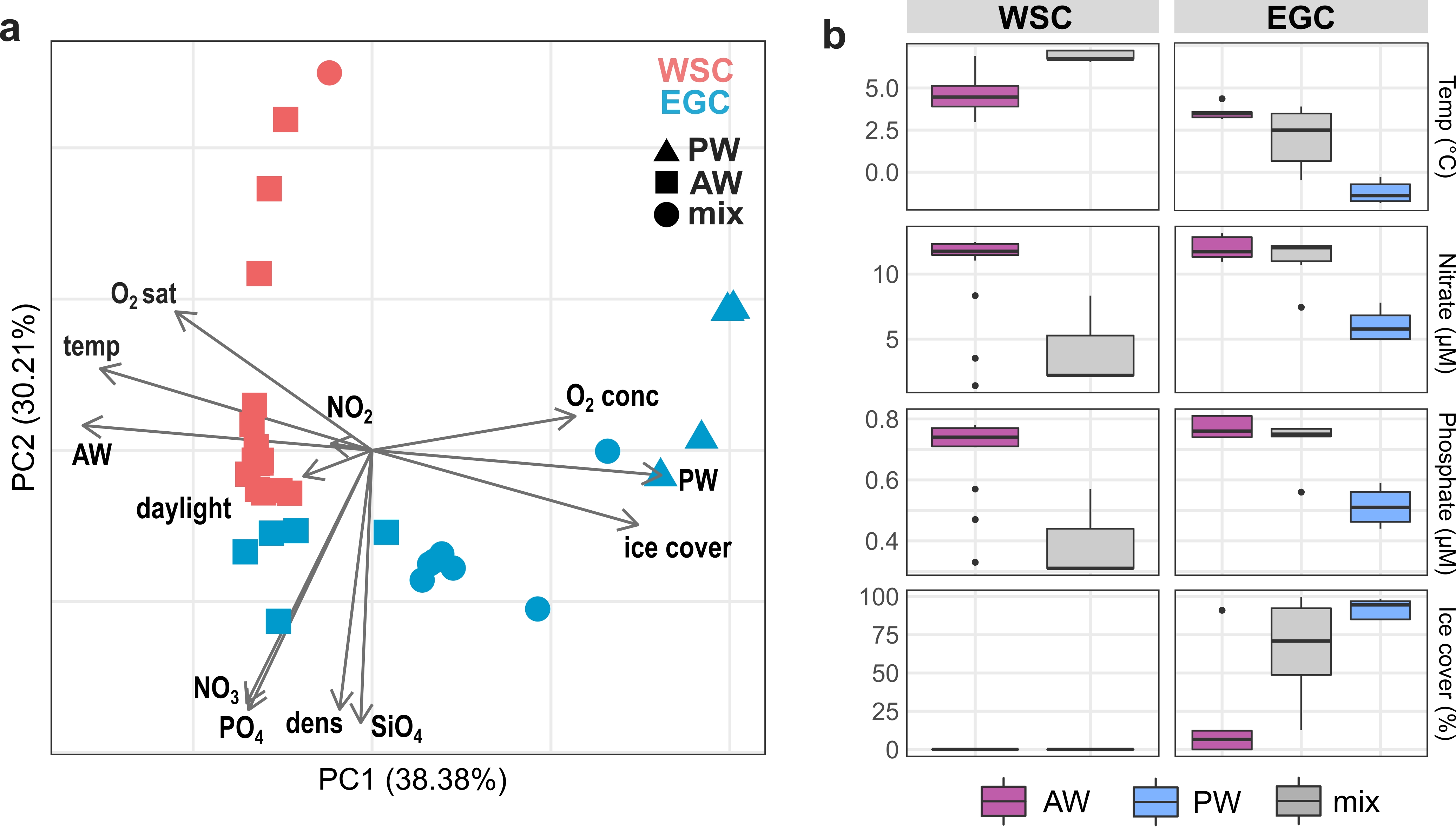

### Extended Data Fig. 2

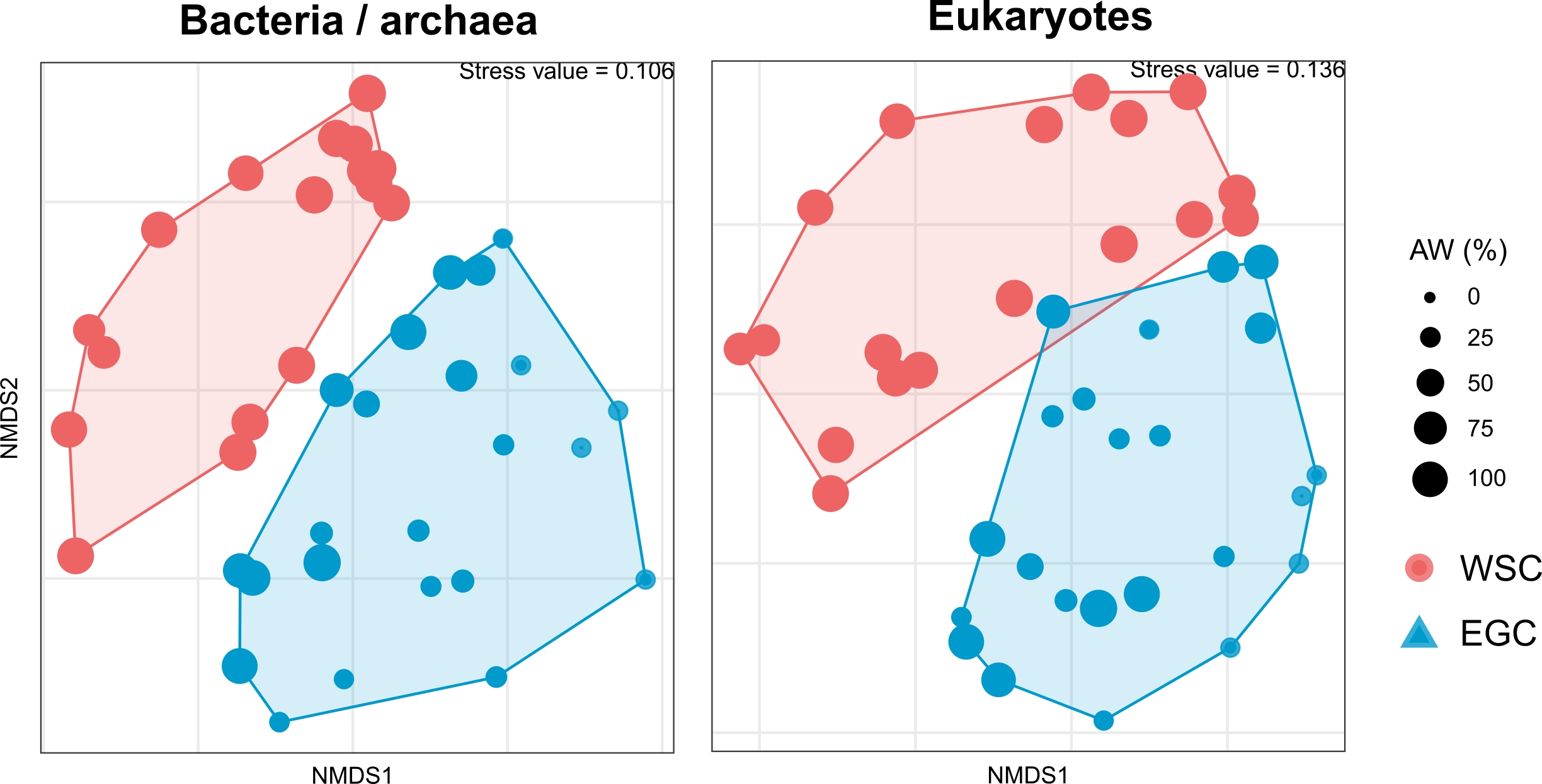

### Extended Data Fig. 3

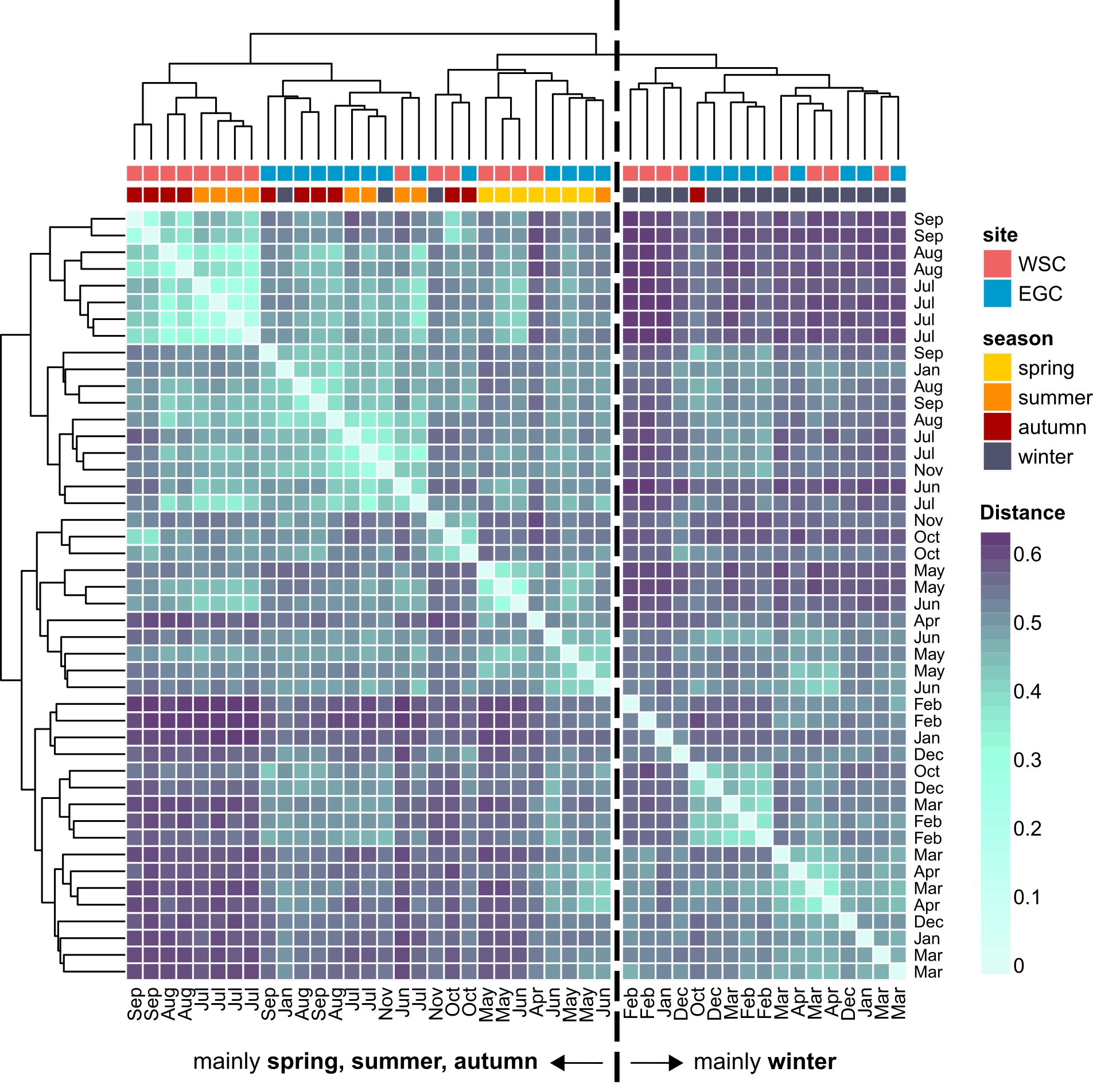

### Extended Data Fig. 4

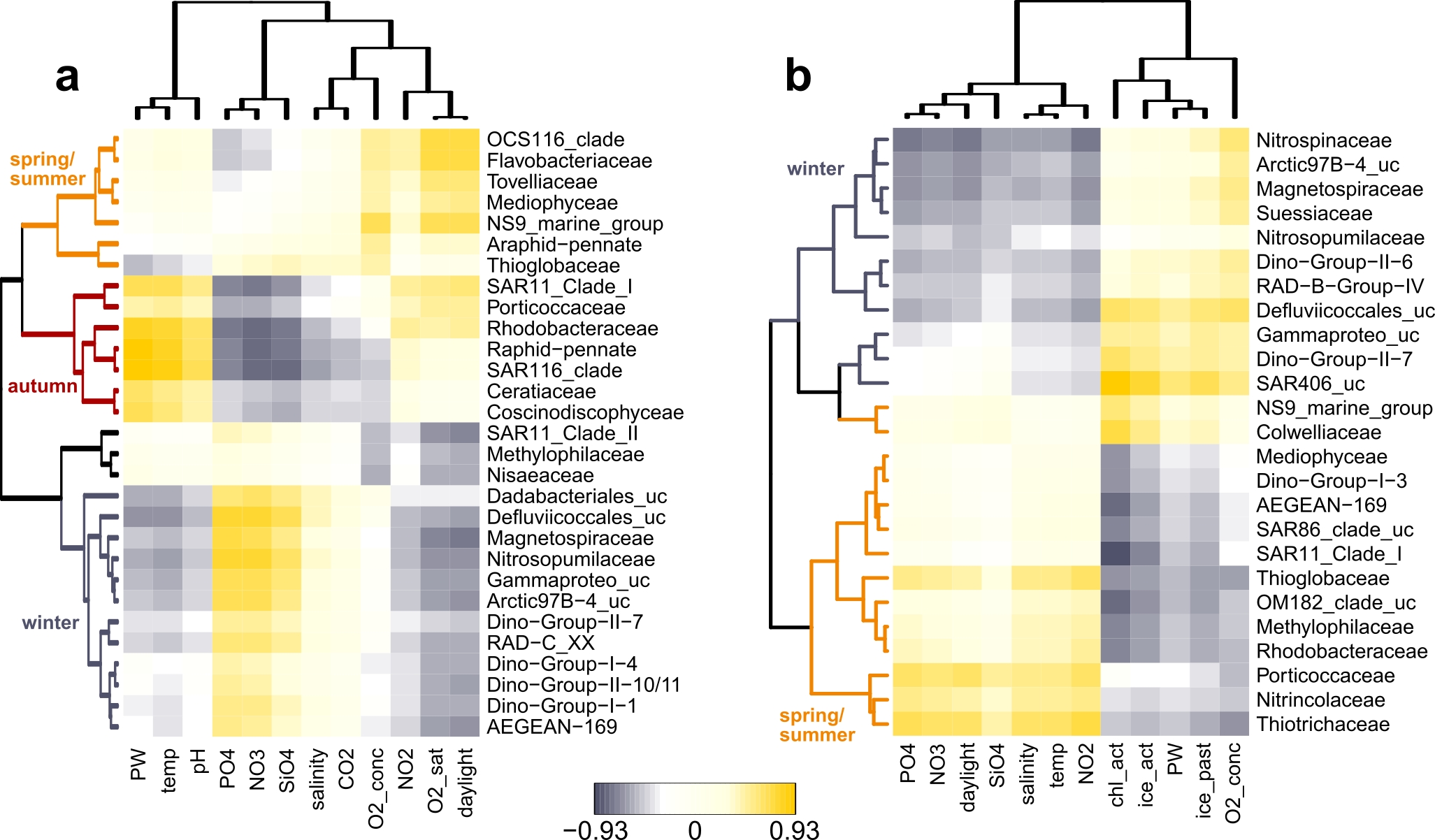

### Extended Data Fig. 5

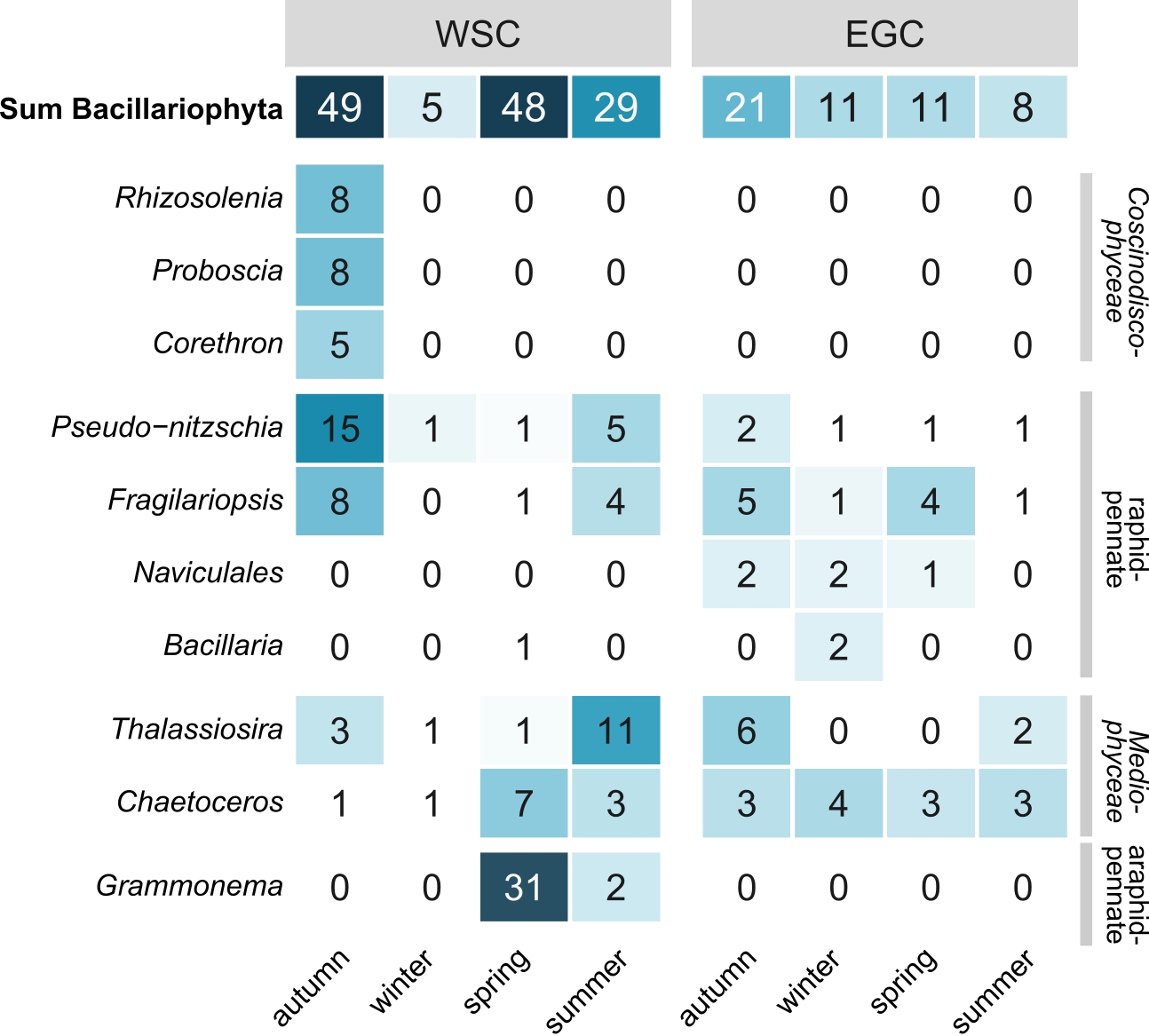

### Extended Data Fig. 6

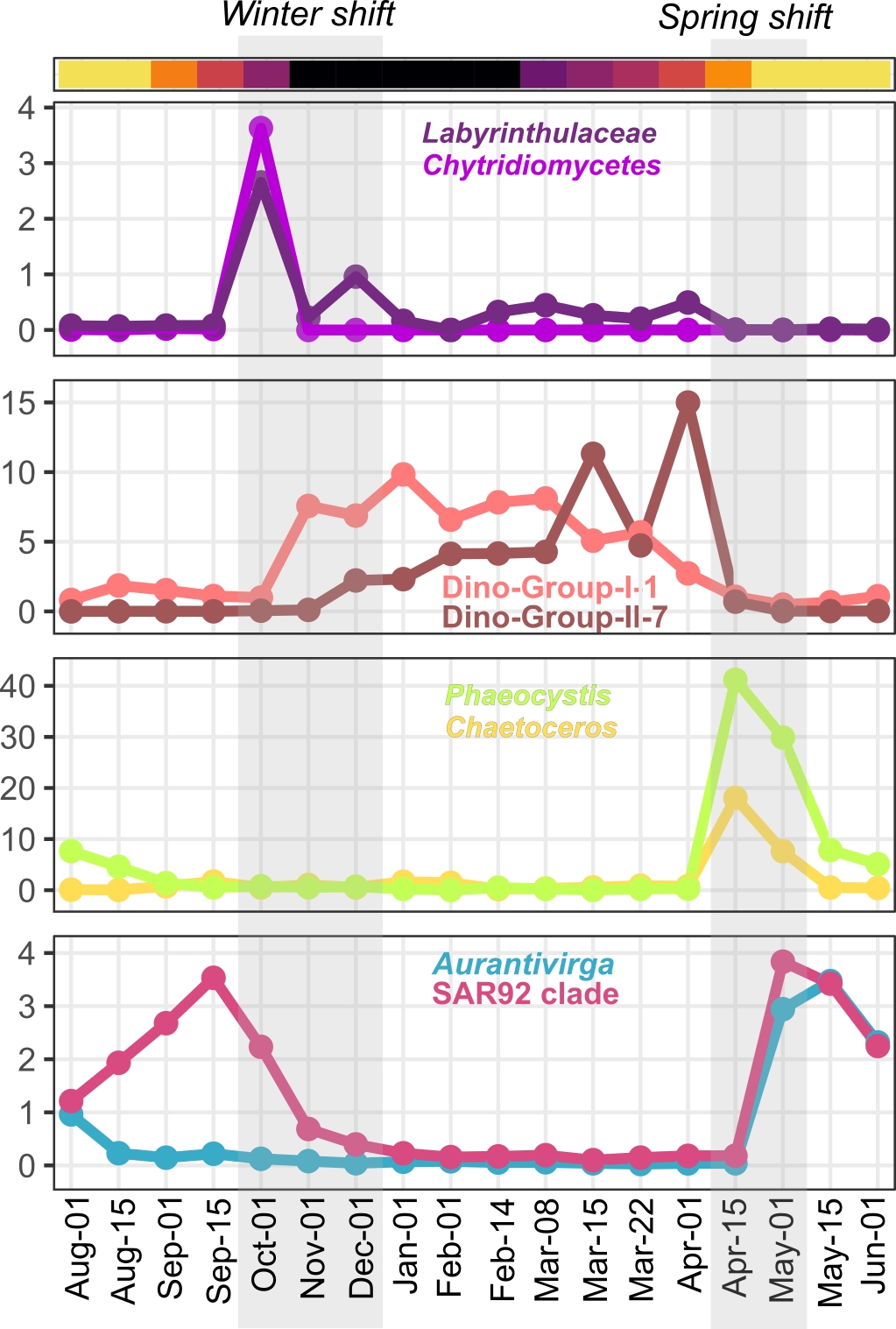

### Extended Data Fig. 7

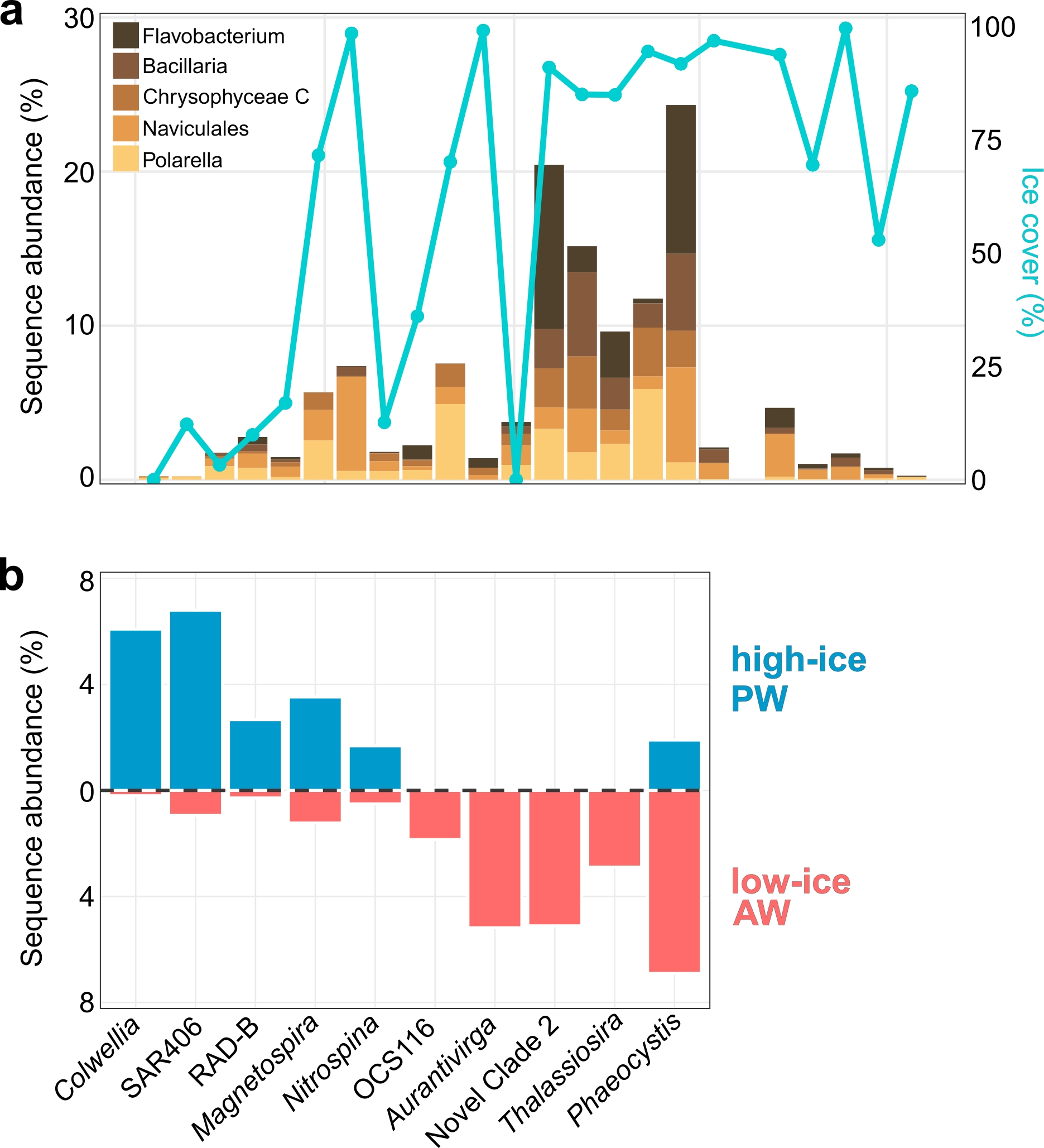

### Extended Data Fig. 8

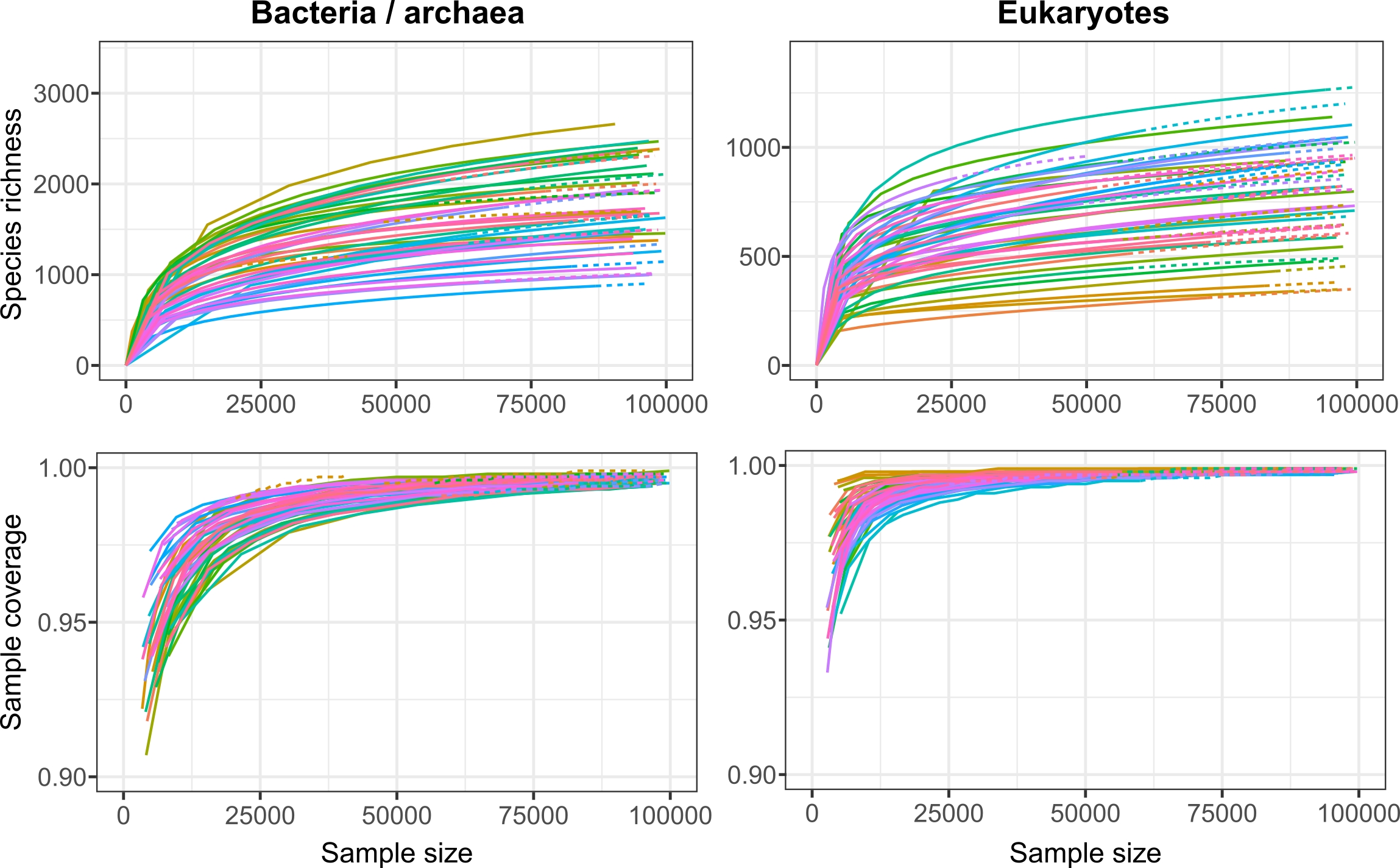
