## Supplementary Material for "The Polar Night Shift: Annual Dynamics and Drivers of Microbial Community Structure in the Arctic Ocean"

### SUPPLEMENTARY FIGURES

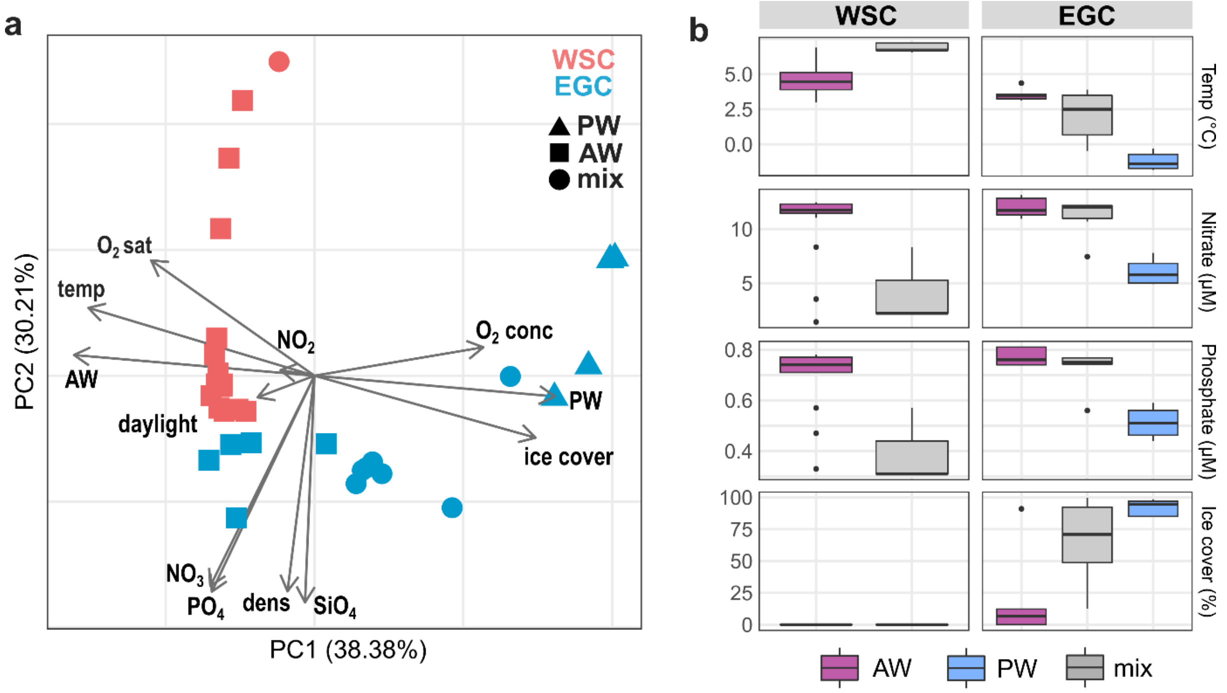


**Extended Data Fig. 1 – Environmental conditions in the Fram Strait. a:** Principal Component Analysis of environmental parameters in the WSC (red) and EGC (blue). Symbol shape designates the prevailing water mass at each sampling event. Arrows illustrate the influence of parameters on multivariate clustering. Temp: water temperature; $O_2$ conc: oxygen concentration; $O_2$ sat: oxygen saturation; AW: proportion of Atlantic Water, PW: proportion of Polar Water; ice: percent ice cover. **b:** Physicochemical parameters with differences between AW, PW and mixed water masses (Kruskal-Wallis test, $p < 0.03$).

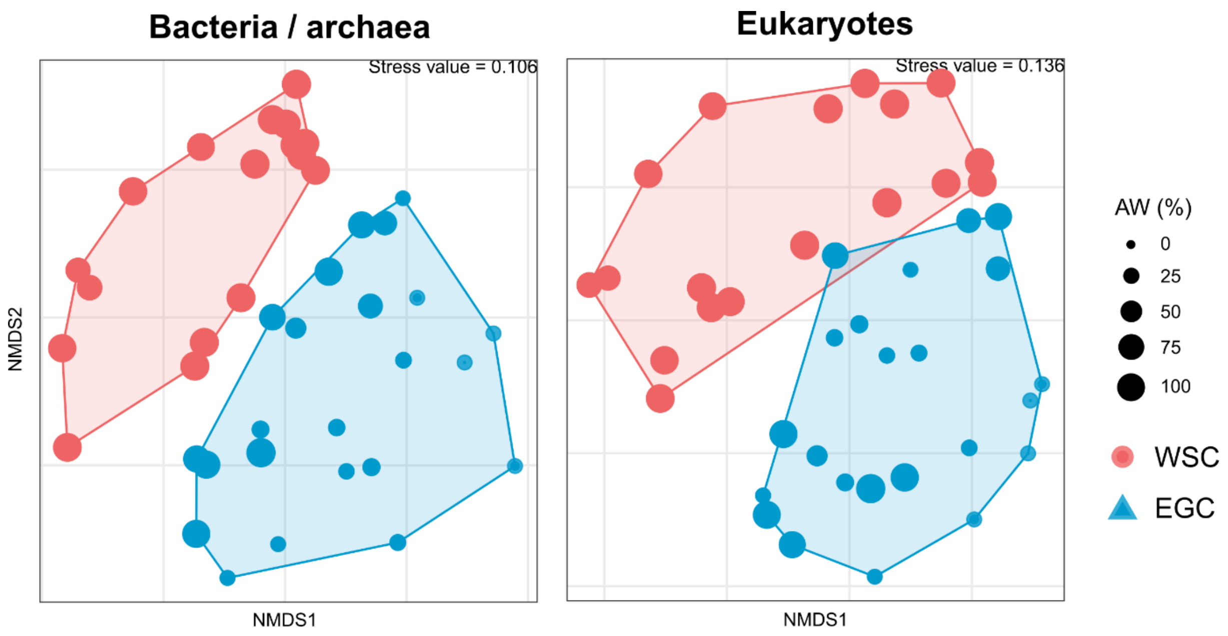


**Extended Data Fig. 2 – Microbial communities in the Fram Strait.** Non-metric multidimensional scaling of Hellinger-transformed relative abundances in the WSC (red) and EGC (blue). Dot sizes correspond to the proportion of Atlantic Water.

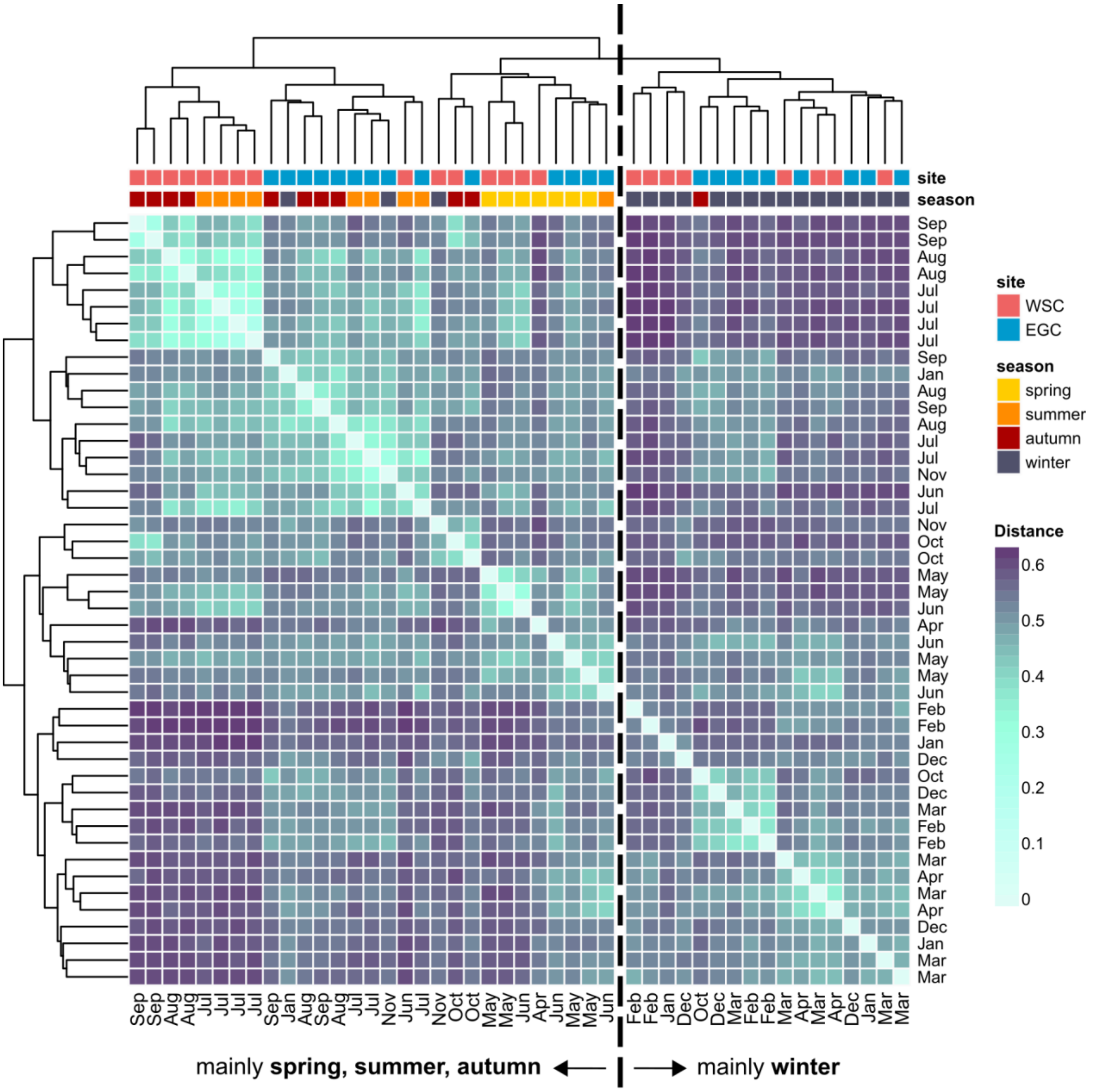


**Extended Data Fig. 3 –** Taxonomic similarities of bacterial and archaeal communities between sampling events and sites expressed as Jenson-Shannon distances.

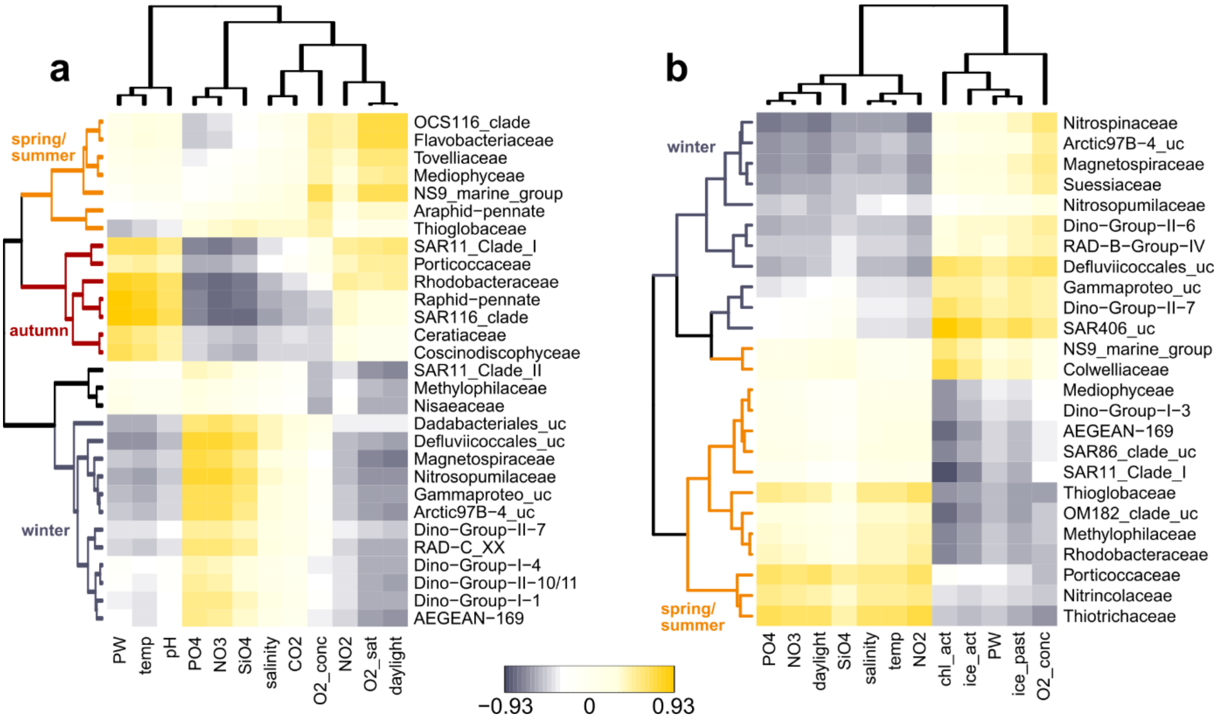


**Extended Data Fig. 4 –** Partial Least Square regression between environmental parameters and the abundance of microbial families. Only correlation coefficients >0.5 were considered. Temp: temperature; $O_2$ conc: oxygen concentration; $O_2$ sat: oxygen saturation; PW: proportion of Polar Water; ice_act: percent ice cover on a given sampling event; ice_past: percent ice cover integrated over the time between sampling events. CO2: partial $CO_2$ pressure

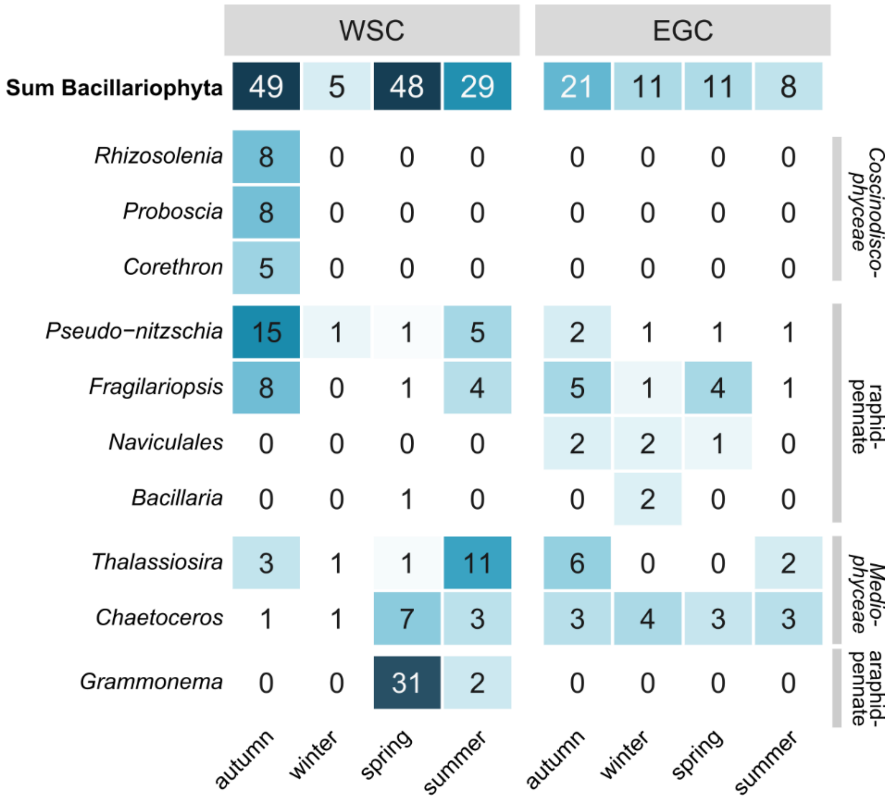


**Extended Data Fig. 5 –** Relative sequence abundances of major diatom genera by season in the WSC and EGC.

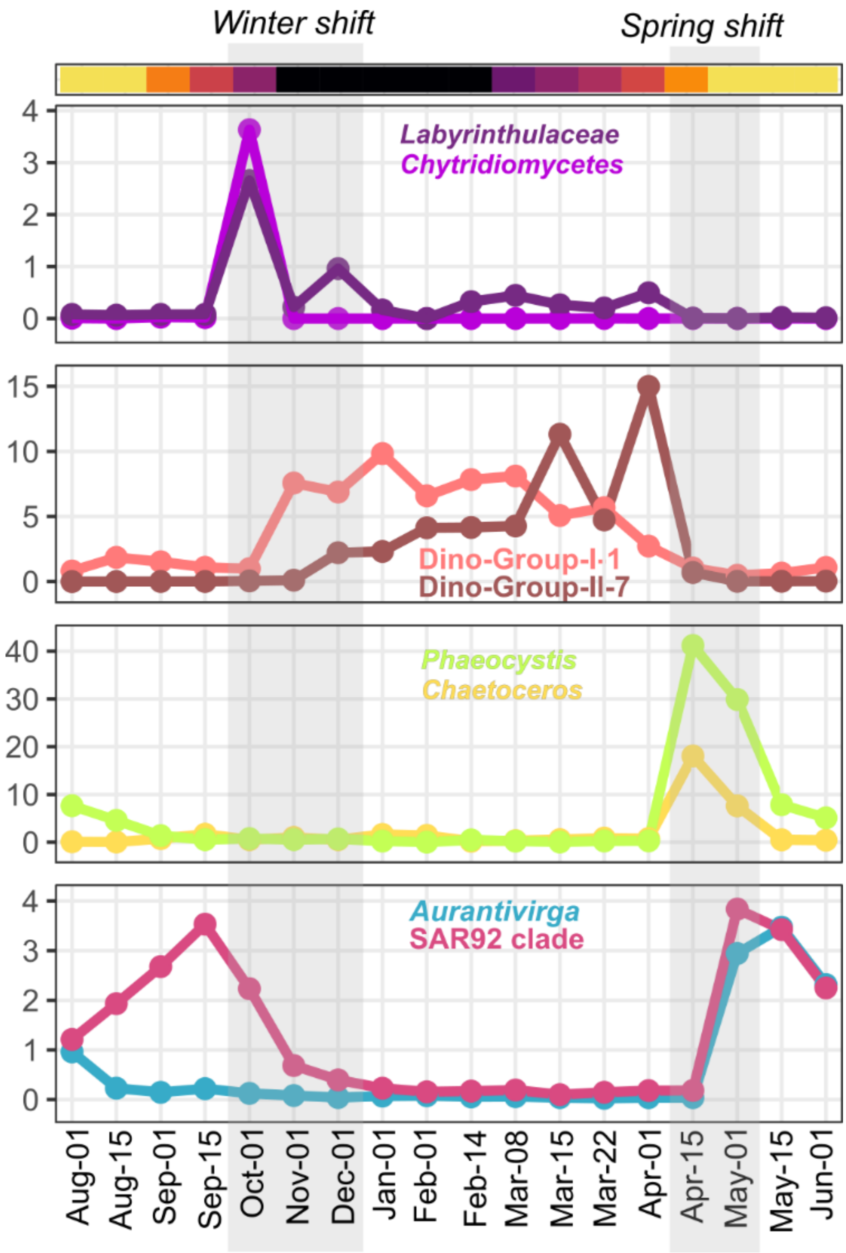


**Extended Data Fig. 6 –** Seasonal transitions in the WSC in relation to daylight hours (color gradient). Late autumn showed a single peak of chytrid fungi and fungal-like *Labyrinthulaceae*, followed by the winter shift to Syndiniales. The winter-to-spring transition featured sudden peaks of the phytoplankton *Phaeocystis* and *Chaetoceros* while Syndiniales disappeared.

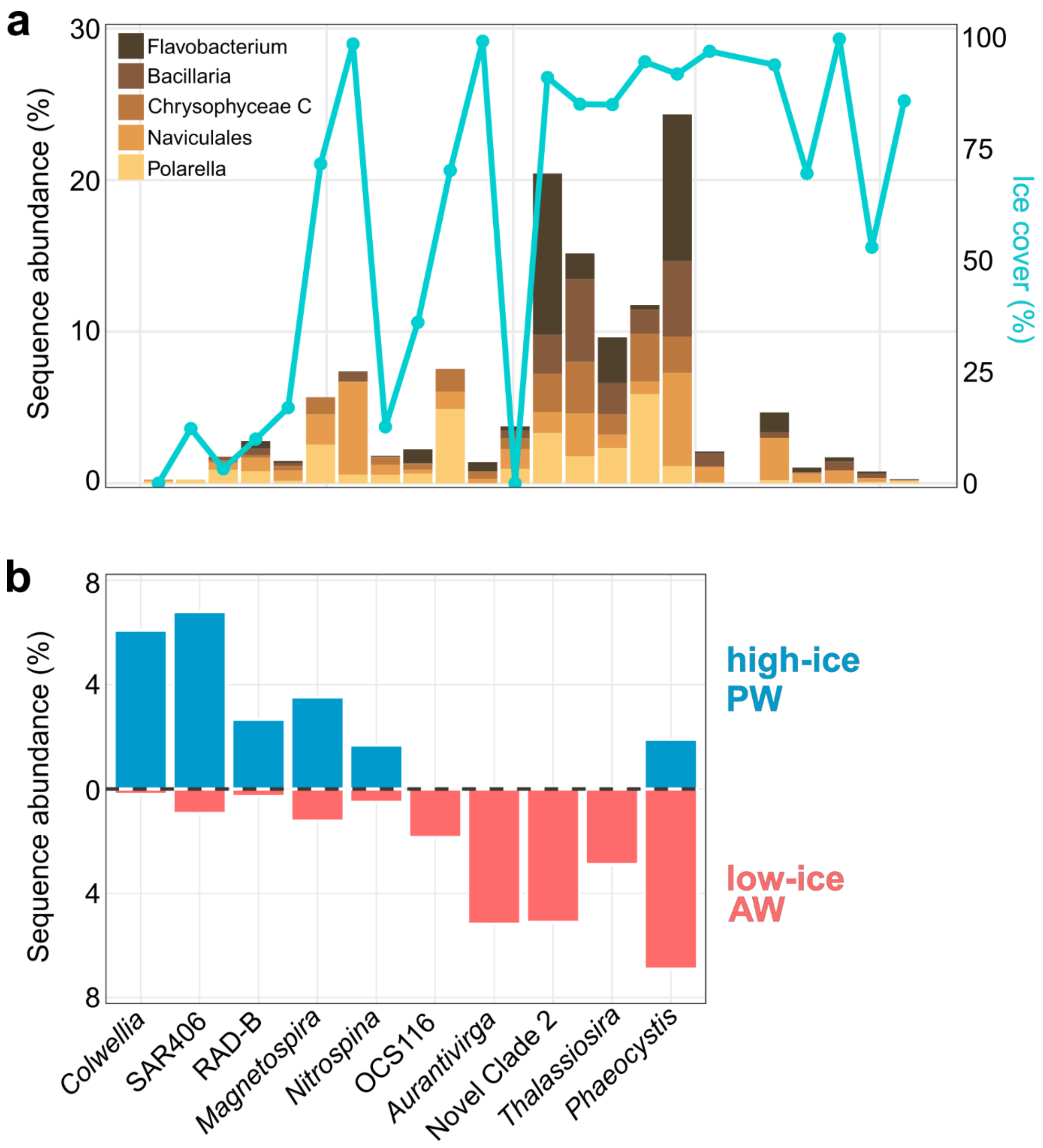


**Extended Data Fig. 7 – EGC-specific patterns. a**: Microbial responses to the ice minimum after intermittent AW advection in January. **b**: Relative sequence abundances of selected genera in polar (high-ice, cold, lower-nutrient) and atlantified (low-ice, warmer, higher-nutrient) conditions.

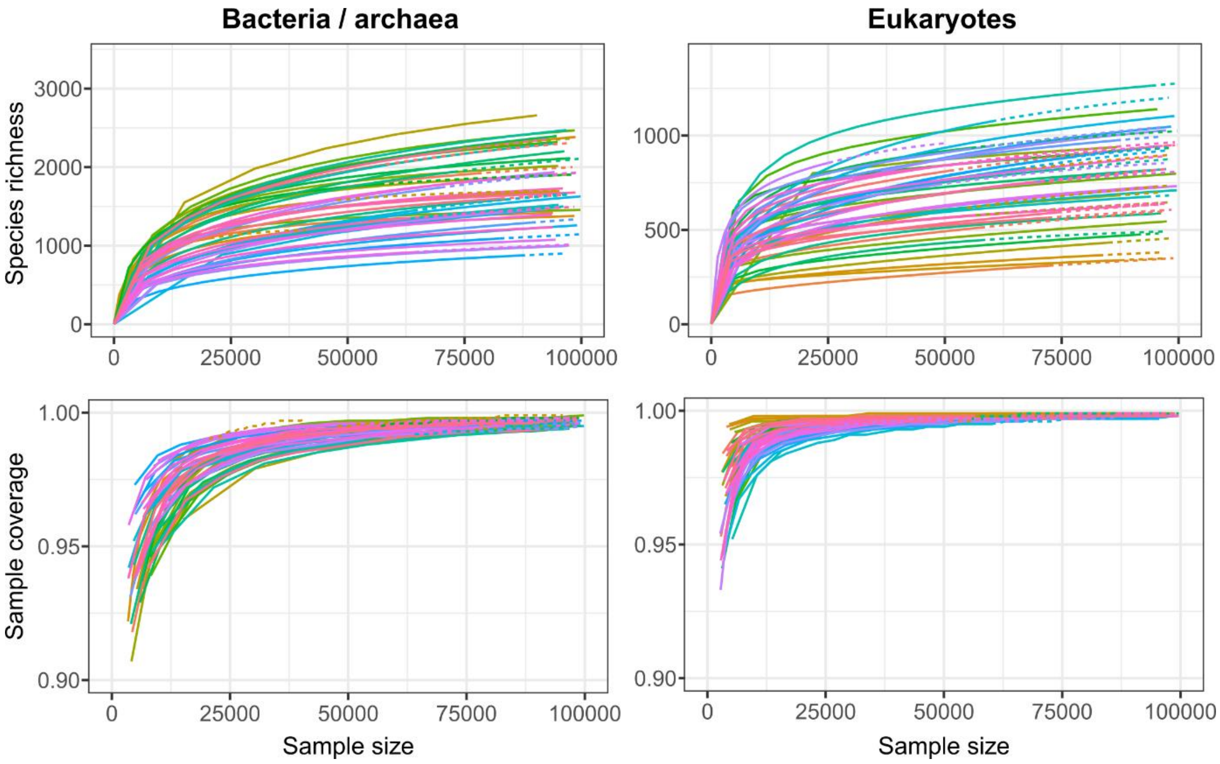


**Extended Data Fig. 8 –** Rarefaction and coverage analyses of amplicon sequence variants, showing that bacterial/archaeal and eukaryotic community composition were sufficiently covered. Each colored line corresponds to an individual sample.

**SUPPLEMENTARY TABLES** (in separate Excel files)

**Supplementary Table 1 –** Overview of sampling events and measured environmental parameters.

**Supplementary Table 2 –** Seasonality of major environmental parameters in the WSC and EGC.

**Supplementary Table 3 –** Amplicon read counts at each major step of the DADA2 pipeline.

### SUPPLEMENTARY METHODS

*Illumina amplicon sequencing*

Library preparation was performed according to the standard instructions of the 16S Metagenomic Sequencing Library Preparation protocol (Illumina, San Diego, CA). The hypervariable V4–V5 region of bacterial 16S rRNA genes was amplified using primers 515F (GTGYCAGCMGCCGCGGTAA) and 926R (CCGYCAATTYMTTTRAGTTT). The hypervariable V4–V5 region of eukaryotic 18S genes was amplified using primers 528iF (GCGGTAATTCCAGCTCCAA) and 926iR (ACTTTCGTTCTTGATYRR). Sequences were obtained on an Illumina MiSeq platform in 2x300 bp paired-end runs at CeBiTec (Bielefeld, Germany) or Alfred Wegener Institute following the standard instructions of the 16S Metagenomic Sequencing Library Preparation protocol (Illumina). Primer-clipped reads were processed into amplicon sequence variants (ASVs) following the standard DADA2 workflow at https://benjjneb.github.io/dada2/tutorial.html[1]. Filtering settings for 16S rRNA amplicons were truncLen = c(230, 195), maxN = 0, minQ = 2, maxEE = c(3, 3) and truncQ = 0, followed by merging using minOverlap = 10 and chimera removal. ASVs were taxonomically classified using the Silva v138 database[2]. Filtering settings for 18S rRNA amplicons were truncLen = c(250, 200), maxN = 0, minQ = 2, maxEE = c(3, 3) and truncQ = 0, followed by merging using minOverlap = 20 and chimera removal. ASVs were taxonomically classified using the PR2 database v4.12[3]. The detailed amplicon workflow has been deposited under https://github.com/matthiaswietz/RAS-1617.

*Characterization of water masses*

Water masses were characterized following[4]. Unlike in that study, only warm Atlantic Water but not Arctic Atlantic Water play a role in our case. As the three types of Atlantic Water (wAW, AW, DW) are temperature-stratified, it is not expected that wAW and DW directly mix without AW in-between. Therefore, two different end member triangles involving only one of wAW and DW can be solved. We define four water masses: warm Atlantic Water (wAW, salinity 35.16, 8°C), Atlantic Water (AW, salinity 35.1, 4.1°C), Polar Surface Water (PSW, salinity

34.17, −1.8°C) and Deep Water (DW, salinity 34.93, −0.9°C). We first applied an end-member decomposition to a wAW-AW-PSW triangle. For some measurements, the solution falls into that triangle meaning that all fractions are between 0 and 1. A number of data points fall to the top/left of the triangle as indicated by negative AW fractions, which we kept as such. A further amount of data points fall to the bottom/right of the triangle, i.e. in the majority into the DW-AW-PSW triangle as indicated by negative wAW fractions. For these cases, we set the wAW fraction to 0 and solve the DW-AW-PSW fractions. This provides four time series for the water masses where at each time point either the wAW or the DW fraction is 0. The total AW fraction (tAW) is defined as the sum wAW+AW+DW. This combines with the PSW time series to add to 1. At times when the water is fresher than PSW, the PSW fraction is > 1 and the tAW is < 0. In those cases, we set the respective values to 1 and 0. Likewise for the cases where tAW > 1. While keeping the details of this calculation in mind, the Polar Water fraction closely follows times of low salinity.
